# Mitochondria remodel near synapses to boost metabolic capacity and fuel plasticity

**DOI:** 10.1101/2025.08.27.672715

**Authors:** Monil Shah, Ilika Ghosh, Nitheyaa Shree Ramesh, Luca Pishos, Tristano Pancani, Valentina Villani, Ryohei Yasuda, Sahil Loomba, Chao Sun, Naomi Kamasawa, Vidhya Rangaraju

## Abstract

Brain synapses are hotspots of energy consumption, and the synaptic plasticity underlying learning sharply increases this demand. Because most synapses lie far from their cell body, their energy demands must be met locally. A dendritic mitochondrion can extend ∼30 μm and span many spines, yet ramps up ATP synthesis only near spines undergoing plasticity, providing immediate and sustained energy. Whether structural changes in these juxtaspinal mitochondrial regions (2 μm segments at spine bases) support this response is unknown; resolving their energy-producing architecture poses substantial challenges. Compared with the lamellar cristae of axonal mitochondria, dendritic mitochondrial cristae form complex, tubular, and interconnected networks that require multiple high-resolution tomograms to characterize. We therefore developed a correlative light and electron microscopy (CLEM) pipeline with a custom deep-learning model that simultaneously segments five distinct 3D suborganellar structures from limited training data, enabling quantitative analysis across 42 tomograms at 2 nm pixel resolution. We find that juxtaspinal mitochondrial regions are larger than non-juxtaspinal regions within the same mitochondrion and enlarge further during synaptic plasticity. Juxtaspinal mitochondrial ATP synthesis and spine ATP levels rise in parallel; during single-spine plasticity, ATP rises within minutes, before detectable enlargement, indicating that juxtaspinal mitochondrial enlargement sustains rather than initiates the local energy response. The enlarged regions exhibit increased cristae membrane surface area, crista junction number, and membrane area with high curvature, together with increased MIC60 and ATP synthase copies, endoplasmic reticulum contacts, and more proximal ribosomes. Matrix-targeted PKA inhibition reduces MIC60 phosphorylation and impairs juxtaspinal mitochondrial enlargement, plasticity-associated ATP increase, and single-spine plasticity. Together, our findings support a model in which spine-specific remodeling within an individual mitochondrion boosts local metabolic capacity to sustain ATP synthesis and fuel synaptic plasticity.

## Introduction

A single dendritic mitochondrion can extend tens of micrometers, spanning dozens of synapses. Organelles of this scale are often studied as single units through measurements of abundance, transport, fusion, and degradation. Yet energy demand within a neuron is spatially heterogeneous, arising at individual synapses, micrometers apart. Whether an organelle can selectively remodel one region to meet local demand, while other regions of the same organelle remain unchanged, remains unresolved.

Neurons pose this problem in its most acute form. Synapses are hotspots of energy consumption, and the plasticity underlying learning is expensive, requiring ATP for ionic homeostasis, protein synthesis, molecular trafficking, and transport. Because most synapses lie hundreds of micrometers from the cell body, where ATP diffusion alone cannot meet these demands, they depend on local supply from spatially stable mitochondria^1,2^. A single dendritic mitochondrion spans many spines yet ramps up ATP production only near those undergoing plasticity, providing immediate (within minutes) and sustained (up to an hour) energy^3^. Whether this functional selectivity reflects a corresponding specialization of mitochondrial structure is unknown. Mitochondria are semi-autonomous organelles carrying their own genome, proteome, and biochemistry, and their inner structure, the organization of cristae and energy-producing complexes, together with contacts to other organelles and the protein synthesis machinery, is a principal determinant of energy output^2-5^. Whether mitochondria actively remodel this inner structure, building energy-producing capacity where and when synapses demand it, remains unknown. This knowledge gap directly impacts human health; disorganized mitochondrial inner structure and interorganellar contacts, with consequent failure to fuel synapses, are hallmarks of many brain disorders^6-8^.

Resolving dendritic mitochondrial structure near synapses presents three technical challenges. First, dendritic mitochondria can extend ∼30 μm and contain complex tubular and network-like cristae, requiring multiple electron tomograms to capture their structural diversity at high resolution. By comparison, short axonal mitochondria (<2 μm), with predominantly lamellar cristae, can be captured more readily within individual tomograms^1,2,5,9^. Second, applying existing deep-learning tools to jointly segment mitochondrial subcompartments and surrounding organelles in plastic-section tomograms requires appropriate training data, adaptation, and validation, complicating automated 3D reconstruction and quantification. Third, linking these ultrastructural measurements to functional readouts, including mitochondrial size changes, ATP levels, and responses to molecular perturbations, requires correlative workflows that identify the same synaptic locations across imaging modalities. These challenges have limited investigations of dendritic, spine-specific mitochondrial adaptations, which remain less characterized than whole-neuron or axonal mitochondrial features^5,10-12^.

To overcome these barriers, we developed a correlative light and electron microscopy (CLEM) pipeline that pairs functional light imaging with electron tomography, together with a custom deep-learning model that simultaneously segments five distinct 3D structures (the volumes enclosed by the outer mitochondrial and inner boundary membranes, the intermembrane space, the endoplasmic reticulum (ER), and ribosomes) from limited training data. Using this pipeline, we examined mitochondrial structure near spines during two forms of synaptic plasticity, homeostatic and single-spine, each of which imposes lasting energy demand^2,3,13-16^. Within a single mitochondrion, 2 μm segments at spine bases (‘juxtaspinal mitochondrial regions’) are larger than equivalent segments without adjacent spines (‘non-juxtaspinal mitochondrial regions’), and enlarge further during plasticity. In intact-brain connectomics datasets^17,18^, juxtaspinal mitochondrial volume scales with the size of large-spine clusters. Juxtaspinal mitochondrial ATP synthesis and spine ATP levels increase during plasticity. Juxtaspinal mitochondrial enlargement accompanies remodeling of the energy-producing architecture, with increased cristae membrane surface area, crista junction number, and membrane area with high curvature, together with increased MIC60 and ATP synthase copies. We also observe increased ER contacts and proximal ribosomes at juxtaspinal mitochondrial regions, suggesting local support for the lipid and protein requirements of remodeling. Matrix-targeted PKA inhibition reduces MIC60 phosphorylation and impairs juxtaspinal mitochondrial enlargement, plasticity-associated ATP increase, and single-spine plasticity. Together, these findings support a model in which a sub-mitochondrial region selectively adapts its inner architecture near individual synapses, boosting local metabolic capacity to sustain ATP synthesis and fuel plasticity.

## Results

### Juxtaspinal mitochondrial regions enlarge upon synaptic plasticity

To examine whether mitochondrial structure changes in response to synaptic plasticity, we labeled neuronal mitochondria with matrix-targeted EGFP (Mito-EGFP) and dendritic spines with mCherry fused to PSD95 (PSD95-mCh) (Fig. 1A, B). To test for spine-specific changes in mitochondrial structure, we measured the area of juxtaspinal mitochondrial regions, 2 μm segments at spine bases (Fig. 1A, see Methods). Dendritic mitochondria are long, often spanning tens of micrometers^1-3,19,20^. So, a single mitochondrion typically extends across juxtaspinal and non-juxtaspinal territory; this let us compare sub-regions within the same mitochondrion, isolating local, intra-mitochondrial changes from whole-organelle differences. Even without plasticity induction, juxtaspinal mitochondrial regions were larger than non-juxtaspinal regions (Fig. 1D).

**Figure 1.**
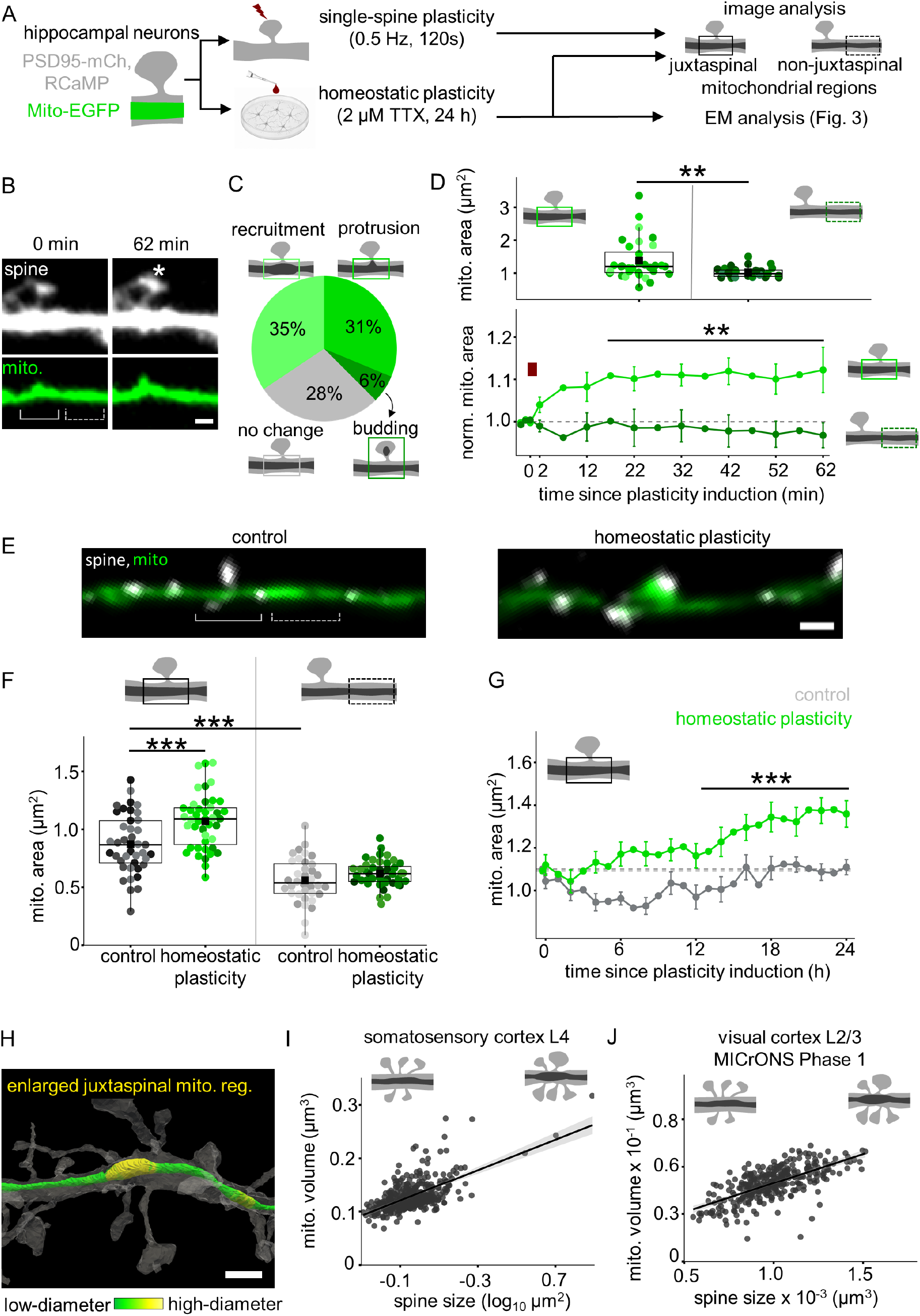
Juxtaspinal mitochondrial regions enlarge upon synaptic plasticity. **A** Schematic of the experimental flow for visualizing juxtaspinal mitochondrial changes by fluorescence and electron microscopy following single-spine and homeostatic plasticity induction. **B** Representative images (quantified in **C, D**) of a dendritic segment upon single-spine plasticity induction, showing spines (white, PSD95-mCh), mitochondria (green, Mito-EGFP), juxtaspinal mitochondrial regions (solid bracket), and non-juxtaspinal mitochondrial regions (dashed bracket). Asterisk, stimulated spine (0.5 Hz, 120 s). Scale bar: 2 μm. **C** Fraction of juxtaspinal mitochondrial regions exhibiting recruitment, protrusion, budding, and no change. n in spines, biol. replicates: 32, 6. **D Top,** baseline average juxtaspinal mitochondrial area is higher compared to non-juxtaspinal mitochondrial area. Different shades within each condition denote independent biological replicates. n in mito. reg., biol. replicates: 32, 6 (juxtaspinal mito. reg.), 30, 6 (non-juxtaspinal mito. reg.). Two-way ANOVA (condition*biol. replicates), p-value: 0.0037. **Bottom,** average juxtaspinal mitochondrial area (green) increases upon single-spine plasticity induction (red bar, 0.5 Hz, 120s); no change in non-juxtaspinal mitochondrial regions (dark green). n in mito. reg., biol. replicates: 32, 6 (juxtaspinal mito. reg.), 30, 6 (non-juxtaspinal mito. reg.). One-way repeated measures ANOVA, Tukey test, p-values: 0.076, 0.006 (juxtaspinal mito. reg.: baseline vs. 2-12 min post-plasticity induction and vs. 17-62 min post-plasticity induction), 0.608, 0.756 (non-juxtaspinal mito. reg.: baseline vs. 2-12 min post-plasticity induction and vs. 17-62 min post-plasticity induction). **E** Representative images (quantified in **F, G**) of a dendritic segment upon homeostatic plasticity induction, showing spines (white, PSD95-mCh), mitochondria (green, Mito-EGFP), juxtaspinal mitochondrial regions (solid bracket), and non-juxtaspinal mitochondrial regions (dashed bracket). Scale bar: 5 μm. **F** Average juxtaspinal mitochondrial area (gray) is greater than the non-juxtaspinal mitochondrial region at control (light gray) and increases further upon homeostatic plasticity induction (green). n in mito.reg., biol. replicates: 41, 3 (juxtaspinal mito. reg. control), 43, 3 (juxtaspinal mito. reg. homeostatic plasticity), 36, 3 (non-juxtaspinal mito. reg. control), 43, 3 (non-juxtaspinal mito. reg. homeostatic plasticity). Three-way ANOVA (condition*mito. region*biol. replicates), Tukey test, p-values: 0.0002 (condition), <0.0001 (juxtaspinal mito. reg.), 0.743 (non-juxtaspinal mito. reg.), <0.0001 (control mito. reg.), 0.0263 (condition*mito. reg.). **G** Average juxtaspinal mitochondrial area increases over 24 hours of homeostatic plasticity relative to control; no change in non-juxtaspinal mito. reg. n in mito.reg., biol. replicates: 90, 3 (juxtaspinal mito. reg. control), 77, 3 (juxtaspinal mito. reg. homeostatic plasticity). One-way repeated measures ANOVA, Tukey test, p-values: 0.422 (juxtaspinal mito. reg., control: baseline vs. 12-24 hrs), <0.0001 (juxtaspinal mito. reg., homeostatic plasticity: baseline vs. 12-24 hrs post-plasticity induction). **H** Representative 3D rendering of a dendritic segment (quantified in **I, J**) showing high- and low-diameter mitochondrial regions within a mitochondrial compartment in the MICrONS Phase 1 connectomics dataset. Color scale indicates local diameter normalized to the mean diameter of the mitochondrial compartment. Scale bar: 1 μm. **I**, **J** Linear fits between local spine size and juxtaspinal mitochondrial volume (within a 2 μm sphere around each spine base) show a significant correlation in the mouse somatosensory cortex (**I**) and visual cortex (**J**). n in synapses, dendrites: 87551, 389 (somatosensory cortex), 767759, 310 (visual cortex). Pearson’s r = 0.63 (somatosensory cortex), 0.66 (visual cortex). p-values: <0.0001 (somatosensory cortex), <0.0001 (visual cortex).

We then induced synaptic plasticity by two paradigms, single-spine and homeostatic (Fig. 1A). For single-spine plasticity, we used two-photon glutamate uncaging to repeatedly stimulate individual spines at 0.5 Hz for 120 s (Fig. 1A-D). Upon induction, juxtaspinal mitochondrial regions enlarged further by one of three routes: recruitment, mitochondrial content increasing; protrusion, mitochondria extending into the plasticity-induced spine; or budding, a mitochondrial fragment pinching off into the spine (Fig. 1C, Supp. video 1; see Methods). Average juxtaspinal mitochondrial area rose from 2 minutes after induction, reached statistical significance at 17 minutes, and remained elevated for up to 62 minutes; no change was detected in non-juxtaspinal mitochondrial regions (Fig. 1D).

Next, we induced homeostatic plasticity by treating primary hippocampal neuronal cultures with 2 μM tetrodotoxin (TTX) for 24 hours and compared them to control neurons (Fig. 1A)^13,15^. We confirmed successful induction from the increased mEPSC amplitude and spine area relative to control, as previously reported^13,14^ (Fig. S1A-C). As above, even without plasticity induction, juxtaspinal mitochondrial regions were larger than non-juxtaspinal regions; after homeostatic plasticity, juxtaspinal mitochondrial regions enlarged further relative to control, whereas non-juxtaspinal regions did not (Fig. 1F). This enlargement appeared as early as ∼12 hours into the 24-hour induction (Fig. 1G). In addition, dendritic, but not axonal, mitochondria became longer relative to control (Fig. S1D, E). These results held when imaging was performed in the continued presence of TTX (Fig. S1F, G, see Methods), and without PSD95 overexpression (using PSD95FingR-EGFP^21^ instead of PSD95-mCh) (Fig. S1H, I).

To determine whether juxtaspinal mitochondrial enlargement also occurs in the intact brain and across cortical areas, we analyzed two volumetric electron microscopy connectomics datasets: mouse visual cortex layer 2/3 (MICrONS Phase 1)^20^ and mouse somatosensory cortex layer 4 ^18^. We developed algorithms to automatically locate spine base centers within dendritic meshes and to measure the juxtaspinal mitochondrial volume around each center (Fig. S1J-L; see Methods). Because these datasets capture static anatomy rather than induced plasticity, we asked whether spine size is associated with local mitochondrial size. In these densely innervated dendrites, a juxtaspinal mitochondrial region was often shared by more than one spine, and non-juxtaspinal mitochondrial regions were uncommon. We, therefore, grouped nearby spines into clusters and related each cluster’s summed spine size to its juxtaspinal mitochondrial volume. Juxtaspinal mitochondrial volume scaled with the summed spine size of each cluster (Fig. 1H-J; see Methods), and these regions displayed morphological signatures of all three enlargement categories (recruitment, protrusion, budding), although in a smaller fraction of spines (Fig. S1M).

These results show that juxtaspinal mitochondrial regions enlarge during synaptic plasticity, across both single-spine and homeostatic plasticity paradigms, with no comparable change detected in non-juxtaspinal or axonal mitochondria. In the intact brain, static anatomy shows a corresponding relationship between juxtaspinal mitochondrial volume and spine size.

### Juxtaspinal mitochondrial regions ramp up ATP synthesis following synaptic plasticity

We next asked whether this spatial asymmetry in structure (Fig. 1B-J, S1G, I) is matched by a spatial asymmetry in energy output, and whether the rise in ATP precedes or follows the structural change. We measured ATP in juxtaspinal and non-juxtaspinal mitochondrial regions (ATP_mito_) and in spines (ATP_spine_) during homeostatic and single-spine plasticity, using our recently developed luciferase-based reporters Mito-ATP and Spn-ATP, respectively^3,22^(Fig. 2A, B, F). Both reporters use an engineered firefly luciferase that emits light by oxidizing luciferin in the presence of ATP and Mg^2+^. Mito-ATP carries four tandem COX8 signal peptides that target it to the mitochondrial matrix (Fig. 2B); Spn-ATP is fused to the postsynaptic scaffold protein Homer2 to target it to spines (Fig. 2F). To control for reporter expression across neurons, conditions, and biological replicates, each reporter is fused to a fluorescent protein (mCherry2 for Mito-ATP and mOrange2 for Spn-ATP), giving a ratiometric luminescence-to-fluorescence (L/F) readout via a custom imaging approach (see Methods)^3,22^. Because both reporters are pH-sensitive, we measured pH independently and applied pH corrections as needed^3^(Fig. S2A, B, D, F-H; see Methods).

**Figure 2.**
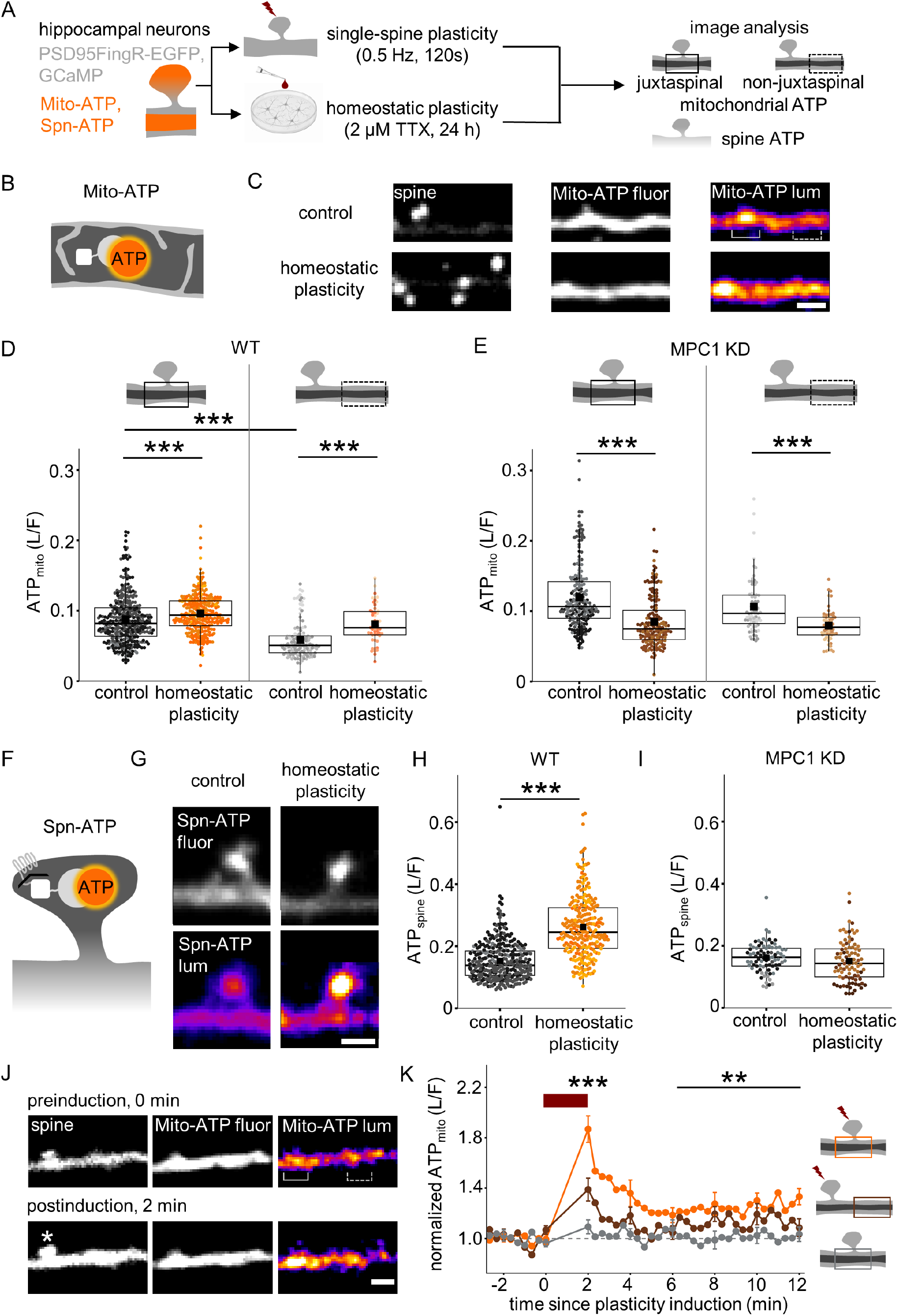
Juxtaspinal mitochondrial regions ramp up ATP synthesis following synaptic plasticity. **A** Schematic of the experimental workflow for imaging ATP_mito_ and ATP_spine_ following single-spine and homeostatic plasticity induction. **B**, **F** Illustrations of the mitochondrial ATP sensor Mito-ATP (**B**), and spine ATP sensor Spn-ATP (**F**). **C** Representative images (quantified in **D**) of a dendritic segment showing spine (white), Mito-ATP fluorescence (white), luminescence (orange), juxtaspinal mitochondrial region (solid bracket), and non-juxtaspinal mitochondrial region (dashed bracket) in control and homeostatic plasticity-induced neurons. Scale bar: 2 μm. **D** Average ATP_mito_ is higher in juxtaspinal than non-juxtaspinal mitochondrial regions under control conditions and increases in both following homeostatic plasticity (orange) relative to control (gray). Different shades within each condition denote independent biological replicates. n in mito. reg., biol. replicates: 352, 3 (juxtaspinal mito. reg. control), 269, 3 (juxtaspinal mito. reg. homeostatic plasticity), 133, 3 (non-juxtaspinal mito. reg. control), 51, 3 (non-juxtaspinal mito. reg. homeostatic plasticity). Three-way ANOVA (condition*mito. reg.*biol. replicates), Tukey test. p-values: <0.0001 (condition), <0.0001 (juxtaspinal mito. reg.), <0.0001 (non-juxtaspinal mito. reg.), <0.0001 (control mito. reg.), 0.1176 (condition*mito. reg.). **E** Upon MPC1 knockdown, average ATP_mito_ decreases in juxtaspinal and non-juxtaspinal mitochondrial regions in homeostatic plasticity-induced neurons (brown) relative to control (gray). n in mito. reg., biol. replicates: 192, 3 (juxtaspinal mito. reg. control), 171, 3 (juxtaspinal mito. reg. homeostatic plasticity), 70, 3 (non-juxtaspinal mito. reg. control), 57, 3 (non-juxtaspinal mito. reg. homeostatic plasticity). Three-way ANOVA (condition*mito. reg.*biol. replicates), Tukey test. <0.0001 (condition), p-values: <0.0001 (juxtaspinal mito. reg.), <0.0001 (non-juxtaspinal mito. reg.), 0.237 (condition*mito. reg.). **G** Representative images (quantified in **H**) showing Spn-ATP fluorescence (white) and luminescence (orange) within a dendritic segment in control and homeostatic plasticity-induced neurons. Scale bar: 2 μm. **H** Average ATP_spine_ increases in homeostatic plasticity-induced neurons (orange) relative to control (gray). n in spines, biol. replicates: 343, 3 (control), 248, 3 (homeostatic plasticity). Two-way ANOVA (condition*biol. replicates), p-value: <0.0001. **I** Upon MPC1 knockdown, average ATP_spine_ fails to increase in homeostatic plasticity-induced neurons (brown) relative to control (gray). n in spines, biol. replicates: 82, 5 (control), 90, 5 (homeostatic plasticity). Two-way ANOVA (condition*biol. replicates), p-value: 0.133. **J** Representative images (quantified in **K**) of a dendritic segment upon single-spine plasticity induction, showing spine (white, GCaMP), Mito-ATP fluorescence (white) and luminescence (orange), juxtaspinal mitochondrial regions (solid bracket), and non-juxtaspinal mitochondrial regions (dashed bracket). Asterisk, plasticity-induced spine (0.5 Hz, 120 s). Scale bar: 2 μm. **K** Average juxtaspinal ATP_mito_ increases upon single-spine plasticity induction (red bar, 0.5 Hz, 120s), with a smaller, non-significant increase in non-juxtaspinal mitochondrial regions and no change in mitochondrial regions near uninduced spines. n in mito. reg., biol. replicates: 25, 11 (juxtaspinal mito. reg.), 16, 10 (non-juxtaspinal mito. reg.), 17, 10, (uninduced). One-way repeated measures ANOVA, Tukey test, p-values: 0.0001, 0.003 (juxtaspinal mito. reg.: baseline vs. 2 min post-plasticity induction and vs. 6-12 min post-plasticity induction), 0.082, 0.04 (non-juxtaspinal mito. reg.: baseline vs. 2 min post-plasticity induction and vs. 6-12 min post-plasticity induction), 0.998, 0.894 (uninduced mito.: baseline vs. 2 min post-plasticity induction and vs. 6-12 min post-plasticity induction).

As with juxtaspinal mitochondrial enlargement, baseline juxtaspinal ATP_mito_ was higher than non-juxtaspinal ATP_mito_ even without plasticity (Fig. 2C, D). After 24 hours of homeostatic plasticity, juxtaspinal ATP_mito_ rose further relative to control (Fig. 2C, D, S2A), on the same timescale as juxtaspinal mitochondrial enlargement, detectable as early as 12 hours (Fig. 1F, G). Non-juxtaspinal ATP_mito_ also rose (Fig. 2C, D), consistent with previous observations of ATP synthesis and distribution across ∼10 μm around plasticity-induced spines^3^. Although the mitochondrial enlargement is confined to juxtaspinal regions (Fig. 1B-G, S1G, I), the ATP it generates extends beyond them to neighboring spines. No such increase in ATP occurred in axonal mitochondria (Fig. S2C, D), consistent with the absence of axonal mitochondrial length changes (Fig. S1E), indicating that this mitochondrial change is specialized to dendritic compartments. The ATP_mito_ increase requires mitochondrial pyruvate entry and downstream ATP synthesis, as it was decreased upon mitochondrial pyruvate carrier knockdown (MPC1 KD) (Fig. 2E). We could not pharmacologically block ATP synthase over the 24-hour timescale of homeostatic plasticity, as the F1F0-ATP synthase inhibitor oligomycin collapsed ATP_mito_ within 10-30 minutes, precluding 24-hour measurements. The rise in ATP_mito_ synthesis led to higher ATP_spine_ relative to control, which was likewise lost upon MPC1 KD (Fig. 2G-I, S2E-G).

To examine single-spine plasticity, we reanalyzed our published dataset^3^, in which individual spines were stimulated by two-photon glutamate uncaging (0.5 Hz, 120s), comparing juxtaspinal mitochondrial regions near plasticity-induced spines with non-juxtaspinal regions and with mitochondrial regions near uninduced spines as controls (Fig. 2J, K). Upon induction, juxtaspinal ATP_mito_ increased within 2 minutes and remained elevated, as previously reported^3^, preceding the mitochondrial enlargement whose mean rose from 2 minutes and reached significance at 17 minutes (Fig. 2J, K, 1D). Non-juxtaspinal ATP_mito_ showed a small, non-significant increase, consistent with both ATP synthesis and distribution across ∼10 μm around plasticity-induced spine^3^ and with our homeostatic plasticity data (Fig. 2J, K, D). Mitochondrial regions near uninduced spines on sister dendrites of the same neurons showed no increase (Fig. 2K). This increase depends on mitochondrial ATP synthesis, as it is abolished by inhibiting F1F0-ATP synthase (with oligomycin) or the mitochondrial pyruvate carrier (with MPC1KD)^3^; these manipulations also block single-spine plasticity^3^.

Together, these results show that juxtaspinal mitochondrial regions are larger and have higher baseline ATP levels than non-juxtaspinal regions, and both rise further with plasticity. Over 24 hours of homeostatic plasticity, the two were not temporally separable, whereas during single-spine plasticity, ATP rose within 2 minutes, preceding detectable mitochondrial enlargement, suggesting that juxtaspinal mitochondrial enlargement supports sustained ATP production rather than the initial ATP burst (see Discussion).

### Juxtaspinal mitochondrial cristae remodel following synaptic plasticity

We next asked whether the inner structure of the juxtaspinal mitochondrial regions changes to expand their ATP production capacity. The inner structure of the mitochondrion, the cristae membrane, crista junctions, and energy-synthesis machinery, is central to energy production^4,5^. Increased cristae membrane density reflects greater energy-production capacity^5,23^. Increased crista junction density reflects tighter association of the cristae with the inner boundary membrane, which is critical for transferring metabolites, including ATP, from the matrix to the intermembrane space and then the cytosol^24^; crista junctions also stabilize the cristae folds and couple to the mitochondrial protein-import machinery^25^. Finally, high cristae curvature reflects local ATP synthase dimerization or multimerization, which drives cristae formation and increases ATP production efficiency in single-celled organisms^26-28^.

Given the specific enlargement of juxtaspinal mitochondrial regions in contrast to non-juxtaspinal regions, we examined cristae only in juxtaspinal mitochondrial regions by establishing a CLEM pipeline (Fig. S3A; see Methods). We used the homeostatic plasticity protocol for CLEM, as it gave substantially higher throughput for identifying and imaging target regions than single-spine plasticity. All measurements were made in dendrites 50-150 μm from the cell body to capture juxtaspinal mitochondrial remodeling in distal segments. Target mitochondria and spines identified by light microscopy were resolved by transmission electron microscopy and imaged with dual-axis tilt series to reconstruct 3D tomograms (21 for control and 21 for homeostatic plasticity; Fig. S3A).

For tomogram analysis, we developed a custom deep-learning model that simultaneously segments five distinct 3D suborganellar structures (the volumes enclosed by the outer mitochondrial and inner boundary membranes, the intermembrane space, the endoplasmic reticulum (ER), and ribosomes) (Fig. S4; see Methods). By learning from multiple tomographic orientations and exploiting spatial context within and between organelles, the model segmented all five structures across 42 tomograms in ∼1 hour (see Discussion).

In correspondence with the enlargement of juxtaspinal mitochondrial regions, juxtaspinal inner mitochondrial membrane (IMM) surface area increased significantly in homeostatic plasticity-induced neurons relative to control (Fig. 3A-D). Separating the cristae membrane from the inner boundary membrane, we quantified juxtaspinal cristae membrane area and crista junction number (Fig. 3A, see Methods). Following homeostatic plasticity, juxtaspinal cristae formed an interconnected, branched network, with increased cristae membrane surface area and more crista junctions relative to control (Fig. 3A-C, E, F, S5, S6, Supp. videos 2-43). We then measured the first-principal-curvature of the cristae membrane, a readout of local ATP synthase dimerization or multimerization^12,29^: homeostatic plasticity increased the juxtaspinal surface area of high-curvature cristae relative to control (Fig. 3A, G, H, S7, 8, Supp. videos 2-43, see Methods).

**Figure 3.**
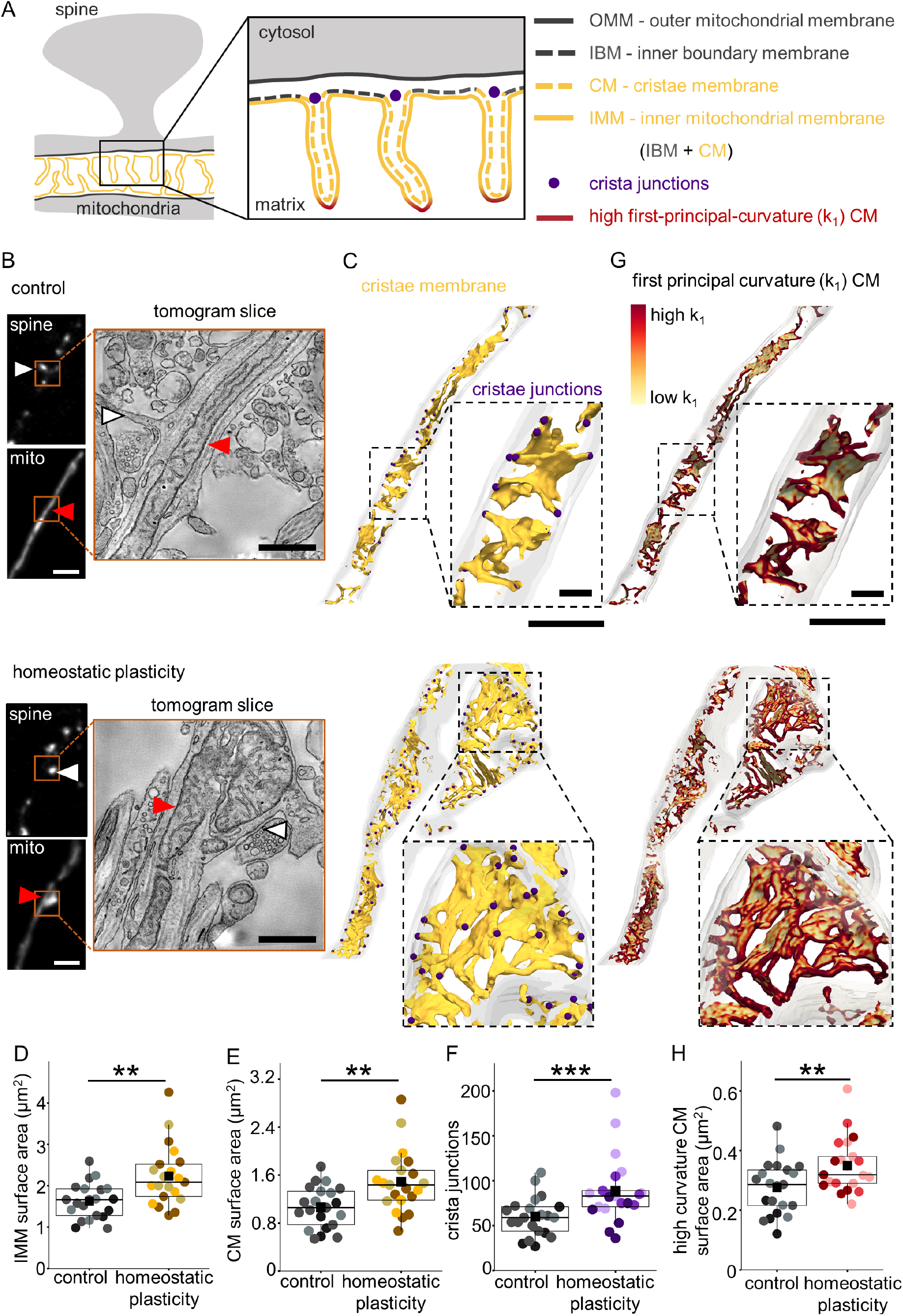
Juxtaspinal mitochondrial cristae remodel following synaptic plasticity. **A** Schematic defining the outer mitochondrial membrane (OMM), inner boundary membrane (IBM), cristae membrane (CM), inner mitochondrial membrane (IMM), crista junctions, and high first-principal-curvature (k_1_) CM used for segmentation and analysis. **B** Representative live confocal images (quantified in **D**-**H**) of a dendritic segment showing spines (white, PSD95-mCh) and juxtaspinal mitochondria (white, Mito-EGFP), subsequently identified and imaged by transmission electron microscopy and tomography (tomogram slice) in control (**top**) and homeostatic plasticity-induced neurons (**bottom**). Scale bars: 2 μm (confocal images), 0.5 μm (EM images). **C** Representative 3D meshes reconstructed from electron tomograms (quantified in **D**-**F,** see also **Fig. S5, 6**) of the juxtaspinal mitochondrial regions boxed in **B** (orange), showing the cristae membrane (yellow) and crista junctions (purple) in control (**top**) and homeostatic plasticity-induced neurons (**bottom**). Scale bar: 0.5 μm. Insets, zoomed-in subregions. Scale bar: 0.1 μm. **D** Average juxtaspinal IMM surface area increases in homeostatic plasticity-induced neurons (yellow) relative to control (gray). Different shades within each condition denote independent biological replicates. n in 3D juxtaspinal mitochondrial reconstructions, biol. replicates: 21, 3 (control), 21, 3 (homeostatic plasticity). Two-way ANOVA (condition*biol. replicates), p-value: 0.0029. **E** Average juxtaspinal cristae membrane surface area increases in homeostatic plasticity-induced neurons (yellow) relative to control (gray). n in 3D juxtaspinal mitochondrial reconstructions, biol. replicates: 21, 3 (control), 21, 3 (homeostatic plasticity). Two-way ANOVA (condition*biol. replicates), p-value: 0.0038. **F** Average number of juxtaspinal crista junctions increases in homeostatic plasticity-induced neurons (purple) relative to control (gray). n in 3D juxtaspinal mitochondrial reconstructions, biol. replicates: 21, 3 (control), 21, 3 (homeostatic plasticity). Two-way ANOVA (condition*biol. replicates), p-value: 0.0009. **G** Representative 3D meshes reconstructed from electron tomograms (quantified in **H**) of the juxtaspinal mitochondrial regions boxed in **B** (orange), showing cristae membrane (yellow) and high-k_1_ cristae membrane (red) in control (**top**) and homeostatic plasticity-induced neurons (**bottom**). Scale bar: 0.5 μm. Insets, zoomed-in subregions. Scale bar: 0.1 μm; color scale spans low k_1_ (0 μm^-1^) to high k_1_ (> 80 μm^-1^). **H** Average juxtaspinal high-k_1_ CM surface area increases in homeostatic plasticity-induced neurons (red) relative to control (gray). n in 3D juxtaspinal mitochondrial reconstructions, biol. replicates: 21, 3 (control), 21, 3 (homeostatic plasticity). Two-way ANOVA (condition*biol. replicates), p-value: 0.0097.

Juxtaspinal mitochondrial enlargement (Fig. 1B-G, S1G, I) is therefore accompanied by increased cristae membrane area, curvature, and junction number, revealing a synaptic plasticity-driven remodeling of the juxtaspinal cristae network consistent with increased ATP-producing capacity and ATP output (Fig. 2C-E, G-K, S2A, B, E-H).

### MIC60 and ATP synthase increase within juxtaspinal mitochondrial regions following synaptic plasticity

MIC60 (mitofilin/IMMT), a core subunit of the MICOS (mitochondrial contact site and cristae organizing system) complex, forms and stabilizes crista junctions, the narrow necks connecting the cristae lumen to the intermembrane space; so a greater number of MIC60-marked junctions reflects a more elaborate cristae network that supports increased metabolite flux, including ATP^24^. In parallel, ATP synthase dimers assemble into rows and higher-order arrays (linear, helical, hexamer, or pyramidal, depending on the species) that impose high membrane curvature and thereby shape the cristae, potentially increasing ATP production capacity^27,30,31^; to date, these arrangements have been resolved almost exclusively in single-celled organisms^4^.

To test whether the juxtaspinal increase in cristae complexity and curvature (Fig. 3B-H, S5-S8) is accompanied by more MIC60 and ATP synthase, we resolved MIC60 and ATP synthase using DNA-PAINT-based single-molecule localization microscopy^32^ (Fig. 4A, B). Across a 60,000-frame acquisition, the cumulative number of localizations per MIC60 copy and per ATP synthase copy reached saturation, confirming complete sampling (Fig. S9A, B), and nearest-neighbor analysis (NeNA) confirmed high localization precision (Fig. S9C, D). We estimated that one copy of MIC60 and one copy of ATP synthase corresponded to 27.76 ± 0.14 and 29.18 ± 0.1 localizations, respectively (Fig. S9E, F).

**Figure 4.**
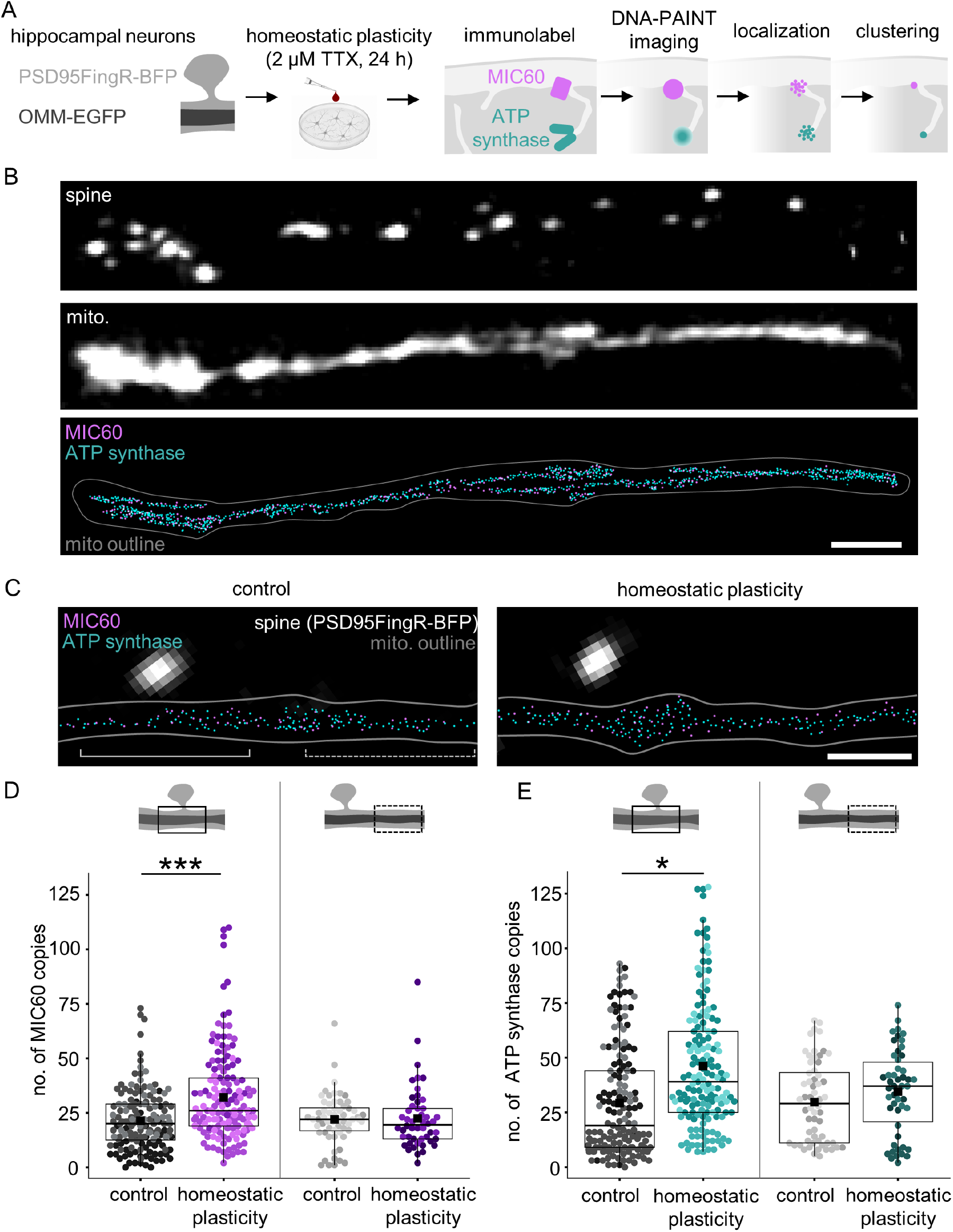
MIC60 and ATP synthase increase within juxtaspinal mitochondrial regions following synaptic plasticity. **A** Schematic of the experimental workflow for visualizing mitochondrial MIC60 and ATP synthase by DNA-PAINT super-resolution microscopy following homeostatic plasticity. **B** Representative images (quantified in **D**, **E**) of a dendritic segment showing spines (**top**; white, PSD95FingR-BFP), mitochondria (**middle**; white, OMM-EGFP), MIC60 (**bottom**; purple, Anti-MIC60), and ATP synthase (**bottom**; turquoise, Anti-OSCP) copies with the mitochondrial boundary overlaid (gray). Scale bar: 2 μm. **C** Representative images (quantified in **D, E**) showing increased MIC60 and ATP synthase copies in juxtaspinal mitochondrial regions following homeostatic plasticity relative to control. Scale bar: 1 μm. **D** Average number of MIC60 copies increases in juxtaspinal mitochondrial regions following homeostatic plasticity (purple) relative to control (gray), but not in non-juxtaspinal mitochondrial regions. Different shades within each condition denote independent biological replicates. n in mito.reg., biol. replicates: 159, 3 (juxtaspinal mito. reg. control), 153, 3 (juxtaspinal mito. reg. homeostatic plasticity), 60, 3 (non-juxtaspinal mito. reg. control), 50, 3 (non-juxtaspinal mito. reg. homeostatic plasticity). Three-way ANOVA (condition*mito. reg.*biol. replicates), Tukey test, p-values: <0.0001 (condition), <0.0001 (juxtaspinal mito. reg.), 0.999 (non-juxtaspinal mito. reg.), <0.0001 (condition*mito. reg.). **E** Average number of ATP synthase copies increases in mitochondrial regions following homeostatic plasticity (turquoise) relative to control (gray). n in mito.reg., biol. replicates: 159, 3 (juxtaspinal mito. reg. control), 153, 3 (juxtaspinal mito. reg. homeostatic plasticity), 60, 3 (non-juxtaspinal mito. reg. control), 50, 3 (non-juxtaspinal mito. reg. homeostatic plasticity). Three-way ANOVA (condition*mito. reg.* biol. replicates), Tukey test, p-values: <0.0001 (condition), 0.027 (juxtaspinal mito. reg.), 0.936 (non-juxtaspinal mito. reg.), 0.117 (condition*mito. reg.).

Following homeostatic plasticity, the detected copy number of MIC60 and ATP synthase per mitochondrion increased (Fig. S9G, H). When resolved by region, this MIC60 increase was confined to juxtaspinal mitochondrial regions, with no change detected in non-juxtaspinal regions (Fig. 4C, D), indicating spine-specific enrichment for MIC60. ATP synthase changed in the same direction, but the regional difference did not reach significance (Fig. 4C, E; see Methods).

These results show that synaptic plasticity increases MIC60 and ATP synthase copy number within enlarged juxtaspinal mitochondrial regions, potentially expanding the local ATP production capacity.

### Juxtaspinal mitochondrial regions gain ER contacts and proximal ribosomes following synaptic plasticity

Mitochondrial proteins are supplied by cytosolic and mitochondrial ribosomes that synthesize proteins on the mitochondrial surface or within the matrix^33,34^. Whole-neuron nascent-proteome profiling after homeostatic plasticity or environmental enrichment detects newly made mitochondrial proteins (both nuclear- and mitochondrially encoded), suggesting that energy demand could drive mitochondrial protein synthesis^13,35^; whether this occurs locally, in distal dendrites and in juxtaspinal mitochondrial regions, is unknown. Expanding the cristae also demands new lipids for the enlarged cristae membrane, sourced from the endoplasmic reticulum (ER) at mitochondria-ER contacts (MERCs)^36-39^.

We first quantified MERCs at juxtaspinal mitochondrial regions in our tomograms (21 control, 21 homeostatic plasticity; Fig. 5A, B, S10, 11, Supp. videos 2-43). Juxtaspinal MERCs increased significantly after homeostatic plasticity relative to control, whereas juxtaspinal ER surface area was unchanged (Fig. 5A-C), indicating more mitochondria-ER contacts rather than simply more ER. Homeostatic plasticity also increased juxtaspinal ribosome number, recruiting ribosomes to within 50 nm of the outer mitochondrial membrane (OMM) (Fig. 5A, B, D, E, S10, 11, Supp. videos 2-43).

**Figure 5.**
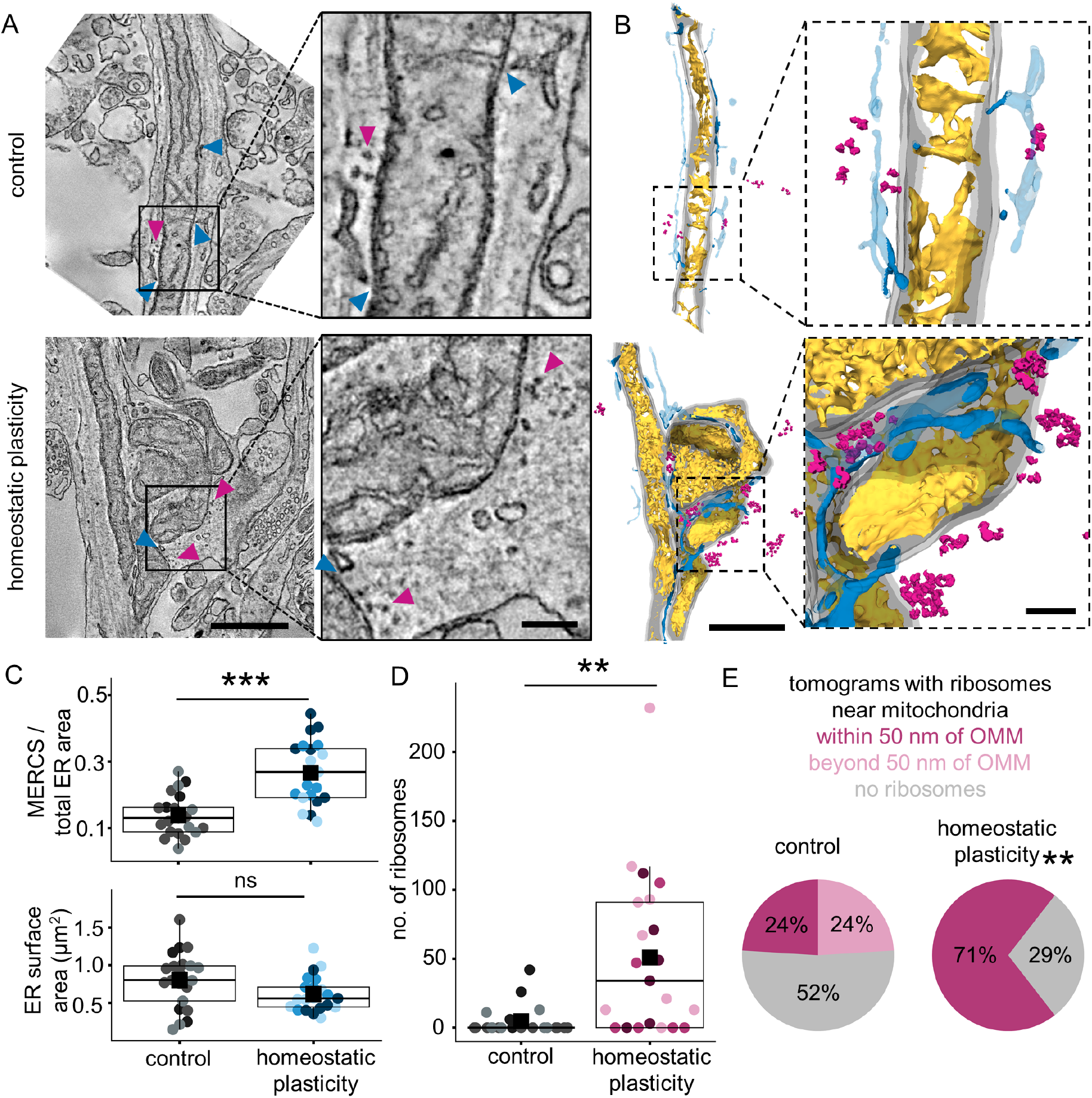
Juxtaspinal mitochondrial regions gain ER contacts and proximal ribosomes following synaptic plasticity. **A** Representative tomogram slices of juxtaspinal mitochondrial regions (quantified in **C**-**E**) identified by the CLEM pipeline in **Fig. S3A** in control (**top**) and homeostatic plasticity-induced neurons (**bottom**), showing mitochondria-ER contacts (MERCS; blue arrowheads), and ribosomes (pink arrowheads). Scale bar: 0.5 μm. Insets, zoomed-in regions. Scale bar: 0.1 μm. **B** Representative 3D meshes reconstructed from the electron microscopy tomograms in **A** (quantified in **C**-**E**), showing MERCS (dark blue), cristae membrane (yellow), ribosomes (pink), and total ER (light blue) in control (**top**) and homeostatic plasticity-induced neurons (**bottom**). Scale bar: 0.5 μm. Insets, zoomed-in regions. Scale bar: 0.1 μm. **C Top,** average fraction of MERCS per total ER increases in homeostatic plasticity-induced neurons (dark blue) relative to control (gray). **Bottom**, average ER surface area is unchanged. Different shades within each condition denote independent biological replicates. n in 3D ER reconstructions around juxtaspinal mito. reg., biol. replicates: 21, 3 (control), 21, 3 (homeostatic plasticity). Two-way ANOVA (condition*biol. replicates), p-values: <0.0001 (top), 0.06 (bottom). **D** Average number of ribosomes near juxtaspinal mitochondrial regions increases in homeostatic plasticity-induced neurons (pink) relative to control (gray). n in 3D ribosome reconstructions around juxtaspinal mito. reg.: 21, 3 (control), 21, 3 (homeostatic plasticity). Two-way ANOVA (condition*biol. replicates), p-value: 0.001. **E** Fraction of ribosomes within 50 nm of the outer mitochondrial membrane (OMM) increases in homeostatic plasticity-induced neurons (pink) relative to control (gray). n in 3D ribosome reconstructions around juxtaspinal mito. reg.: 21, 3 (control), 21, 3 (homeostatic plasticity). Fisher’s exact test, p-value: 0.0048.

Thus, alongside cristae remodeling, synaptic plasticity locally increases MERCs and mitochondria-proximal ribosomes in juxtaspinal mitochondrial regions, positioning the machinery for lipid transfer and protein synthesis to supply the building blocks for the remodeling.

### Juxtaspinal mitochondrial remodeling requires matrix PKA activity and MIC60 phosphorylation to fuel plasticity

To determine whether juxtaspinal mitochondrial remodeling is required to sustain ATP synthesis and fuel synaptic plasticity, we sought a molecular handle to block the remodeling. Many proteins shape cristae, such as OPA1, tafazzin, the MICOS subunits (MIC10, MIC19, MIC60), prohibitins, stomatin-like protein-2, and DNAJC19, but genetic deletion of any of them grossly disorganizes cristae and compromises mitochondrial function and cell health^40-44^, and most work has been in isolated mitochondria, yeast, or non-neuronal cells. However, which player(s) drive juxtaspinal mitochondrial remodeling after synaptic plasticity is unknown. PKA, a cAMP-dependent kinase, phosphorylates MIC60 and MIC19 in non-neuronal mammalian cells^7,45,46^. Because PKA is a signaling kinase rather than a structural component of the cristae machinery, we reasoned that inhibiting PKA could block remodeling without dismantling the cristae architecture. Moreover, because matrix cAMP is generated by soluble adenylyl cyclase in response to metabolic state, intramitochondrial cAMP-PKA signaling^47-49^ is well positioned to couple cristae remodeling to synaptic energy demand.

Because PKA is ubiquitous, we targeted its inhibition to mitochondrial compartments. We genetically targeted PKI, a heat-stable 76-amino-acid peptide inhibitor selective for the PKA catalytic subunit^50^, to either the mitochondrial matrix (Matrix-PKI) or the intermembrane space (IMS-PKI) (Fig. 6A). We then used MIC60 phosphorylation as a readout^49^ to assess the effects of compartment-targeted PKA inhibition.

**Figure 6.**
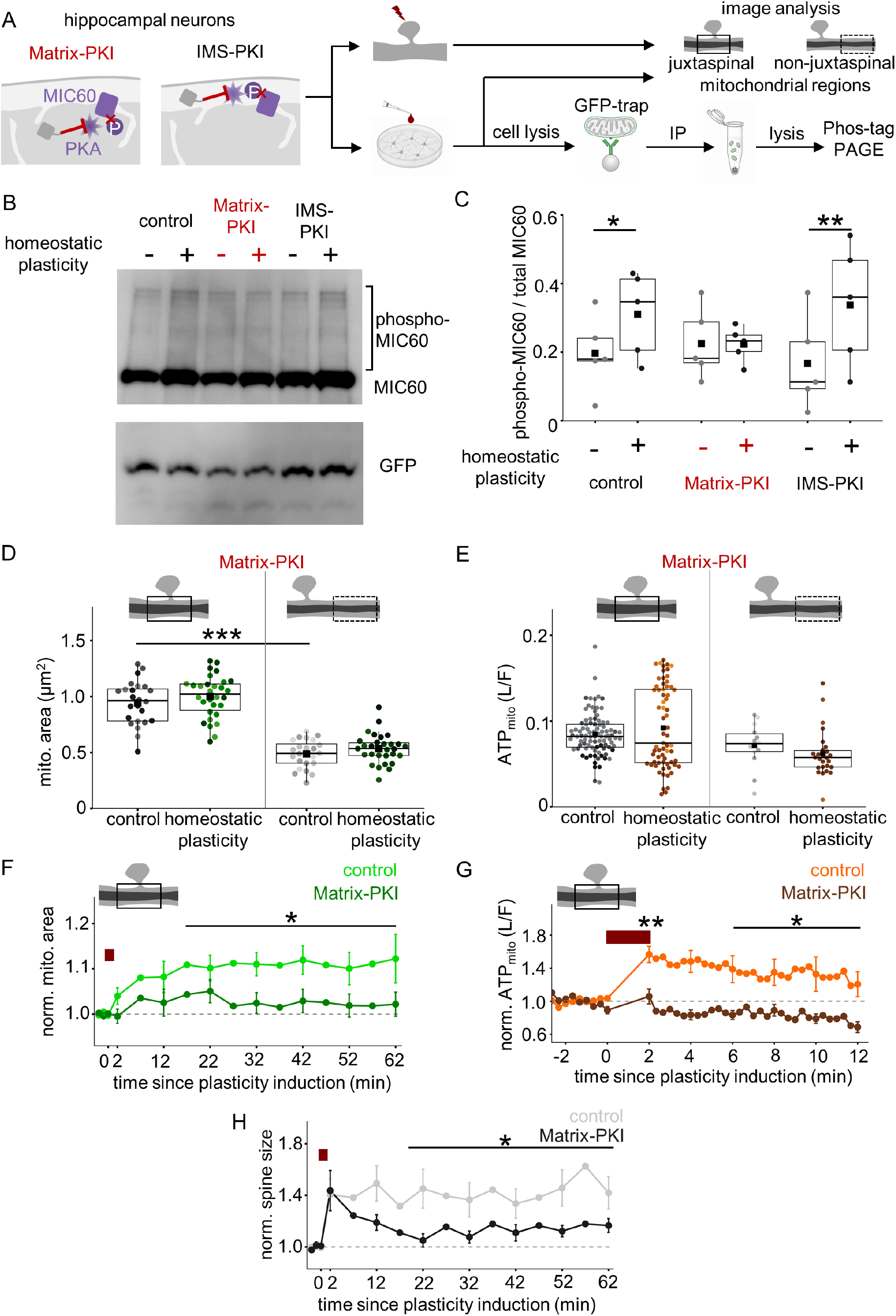
Juxtaspinal mitochondrial remodeling requires matrix PKA activity and MIC60 phosphorylation to fuel synaptic plasticity. **A** Schematic of the experimental workflow for PKA inhibition using Matrix-PKI and IMS-PKI during single-spine and homeostatic plasticity induction. **B** Representative Phos-tag mobility shift blot following homeostatic plasticity induction in control (black), Matrix-PKI (red), and IMS-PKI (black) expressing neurons. Mitochondria were isolated by OMM-GFP pulldown; OMM-GFP served as the loading control and to confirm immunoprecipitation. **C** Average ratio of MIC60 phosphorylation-to-total MIC60 signal showing increased MIC60 phosphorylation following homeostatic plasticity in control and IMS-PKI-expressing neurons, but not in Matrix-PKI-expressing neurons. Different shades within each condition denote independent biological replicates. n in blots, biol. replicates: 5, 5 (control, +/-homeostatic plasticity), 5, 5 (Matrix-PKI, +/-homeostatic plasticity), 5, 5 (IMS-PKI, +/-homeostatic plasticity). Two-way (condition*treatment) matched ANOVA, Tukey test, p-values: 0.0407 (control), 0.967 (Matrix-PKI), 0.0046 (IMS-PKI). **D** In Matrix-PKI expressing neurons, average juxtaspinal mitochondrial area remains greater than non-juxtaspinal mitochondrial regions under control conditions (gray) but fails to increase further upon homeostatic plasticity induction (green). n in mito. reg., biol. replicates: 24, 3 (juxtaspinal mito. reg. control), 30, 3 (juxtaspinal mito. reg. homeostatic plasticity), 24, 3 (non-juxtaspinal mito. reg. control), 30, 3 (non-juxtaspinal mito. reg. homeostatic plasticity). Three-way ANOVA (condition*mito. reg.*biol. replicates), Tukey test, p-values: 0.092 (condition), <0.0001 (control mito. reg.), 0.9 (condition*mito. reg.). **E**. **I**n Matrix-PKI expressing neurons, average ATP_mito_ is no longer higher in juxtaspinal than non-juxtaspinal mitochondrial regions under control conditions, and fails to increase following homeostatic plasticity (brown) relative to control (gray). n in mito.reg., biol. replicates: 91, 3 (juxtaspinal mito. reg. control), 67, 3 (juxtaspinal mito. reg. homeostatic plasticity), 14, 3 (non-juxtaspinal mito. reg. control), 26, 3 (non-juxtaspinal mito. reg. homeostatic plasticity). Three-way ANOVA (condition*mito region*biol. replicates), Tukey test, p-values: 0.871 (condition), 0.970 (control mito. reg.), 0.338 (condition*mito. reg.). **F** In Matrix-PKI expressing neurons (dark green), average juxtaspinal mitochondrial area fails to increase upon single-spine plasticity induction (red bar, 0.5 Hz, 120s) relative to control (green). n in juxtaspinal mito. reg., biol. replicates: 32, 6 (control), 22, 3 (Matrix-PKI). Kruskal-Wallis ANOVA test, p-values: 0.0303 (norm. juxtaspinal mito. reg. 17-62 min post-plasticity induction: control vs. Matrix-PKI). **G** In Matrix-PKI-expressing neurons (orange), average juxtaspinal ATP_mito_ fails to increase upon single-spine plasticity induction (red bar, 0.5 Hz, 120s) relative to control (brown). n in mito. reg., biol. replicates: 13, 3 (control), 9, 3 (Matrix-PKI). One-way ANOVA, Tukey test, p-values: 0.0016, 0.013 (norm. juxtaspinal mito. reg. 2 min and 6-12 min post-plasticity induction: control vs. Matrix-PKI). **H** In Matrix-PKI-expressing neurons (black), average spine size fails to show sustained increase upon single-spine plasticity induction (red bar, 0.5 Hz, 120s) relative to control (gray). n in spines, biol. replicates: 32, 6 (control), 22, 3 (Matrix-PKI). Two-way repeated measures ANOVA (condition*time), p-values: 0.0232 (17-62 min post-plasticity induction: control vs. Matrix-PKI).

**Figure 7.**
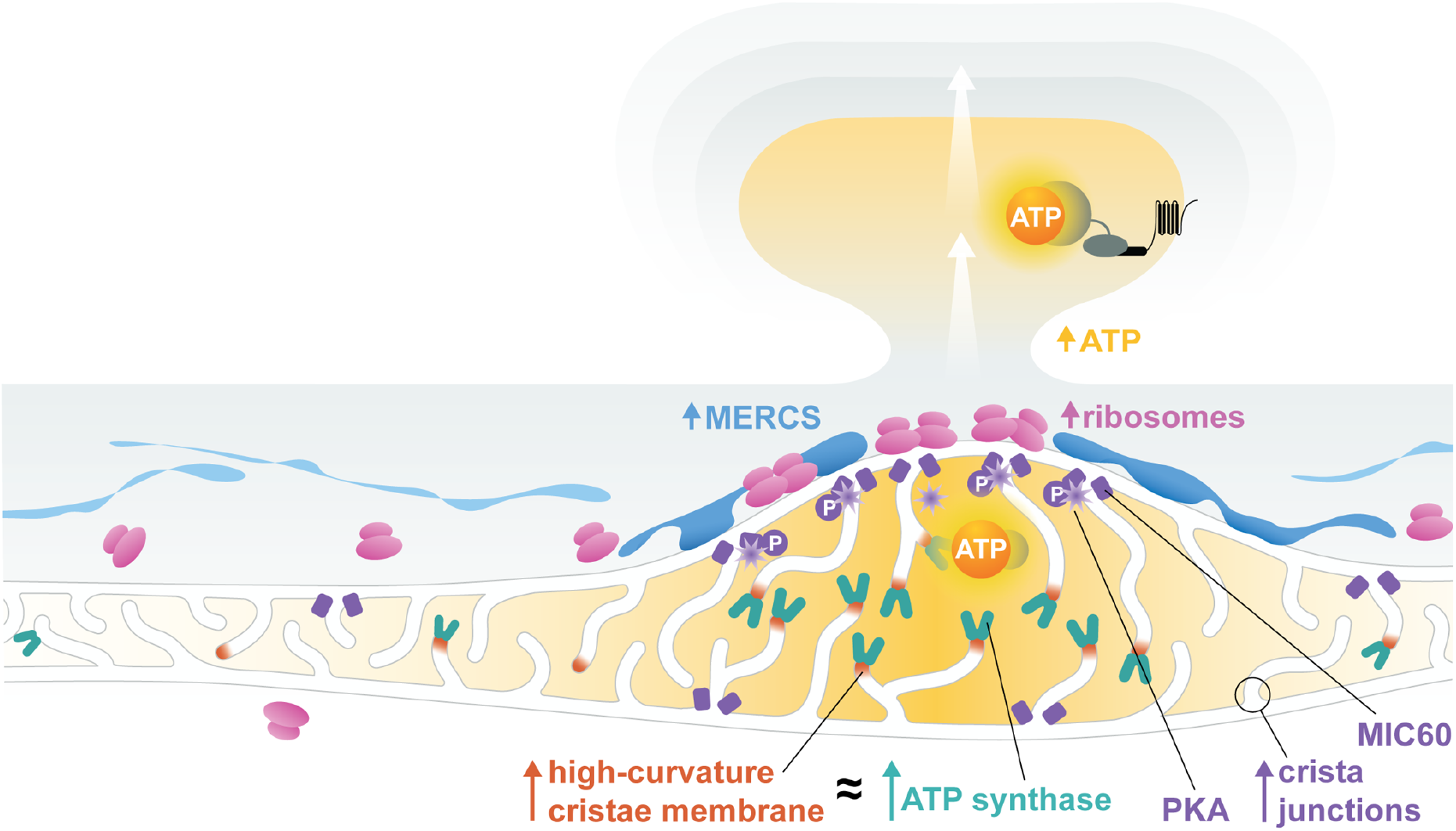
Mitochondria remodel near synapses to boost metabolic capacity and fuel plasticity. Summary schematic of juxtaspinal mitochondrial remodeling upon synaptic plasticity. Remodeling comprises increases in cristae membrane (gray line), crista junctions (black circle), high cristae curvature (red), MIC60 (purple), ATP synthase (turquoise), mitochondria-endoplasmic reticulum contacts (MERCS, dark blue; ER, light blue), and ribosomes near mitochondria (pink); together, these expand local energy-producing capacity to sustain mitochondrial ATP synthesis and spine ATP levels (orange). Remodeling requires matrix PKA activity (purple star) and MIC60 phosphorylation (purple P-tag). Mitochondrial ultrastructure, ER contacts, and ribosomes were measured using the CLEM pipeline with deep-learning-based suborganellar segmentation and 3D reconstruction.

Homeostatic plasticity increased MIC60 phosphorylation, detected by Phos-tag mobility shift (Fig. 6B, C). This increase was abolished by Matrix-PKI expression, but not by IMS-PKI expression, supporting a role for matrix PKA activity in regulating MIC60 phosphorylation (Fig. 6B, C). Homeostatic plasticity also increased phosphorylation of the cytosolic PKA substrates GluA1 and GluN1, consistent with previous reports^51-53^, but neither Matrix-nor IMS-PKI altered these responses (Fig. S12A-D), supporting compartment-selective inhibition.

We then asked whether blocking mitochondrial matrix PKA affects the plasticity response. Matrix-PKI failed to increase juxtaspinal mitochondrial area and ATP_mito_ synthesis (Fig. 6D, E, S12E). Baseline measurements were unaffected: Matrix-PKI did not alter resting mitochondrial area, juxtaspinal mitochondrial regions remained wider than non-juxtaspinal mitochondria (Fig. 6D compared to Fig. 1F, p-value: 0.243); Matrix-PKI did not alter baseline ATP_mito_ (Fig. 6E compared to Fig. 2D, p-value: 0.489), indicating that the effect of matrix PKA is specifically engaged in response to plasticity rather than acting tonically (see Discussion). Notably, homeostatic plasticity itself, measured as increased mEPSC amplitude, was preserved (Fig. S12F), suggesting that alternative fuel sources such as glycolysis might compensate for the lost mitochondrial ATP increase over the 24-hour induction. By contrast, single-spine plasticity, an acute local demand, was abolished by Matrix-PKI along with juxtaspinal mitochondrial enlargement and ATP_mito_ synthesis (Fig. 6F-H, S12G), consistent with a requirement for local mitochondrial ATP that glycolysis cannot meet on this timescale^3^. Therefore, matrix PKA activity, accompanied by PKA-dependent MIC60 phosphorylation, is required for juxtaspinal mitochondrial enlargement, ramping up local mitochondrial ATP synthesis and supporting synaptic plasticity.

Overall, our findings support the model that juxtaspinal mitochondrial remodeling is a spine-specific, sub-organellar specialization that expands mitochondrial metabolic capacity to sustain ATP production and fuel synaptic plasticity.

## Discussion

We show that dendritic mitochondria remodel their inner structure within juxtaspinal mitochondrial regions, expanding their capacity to sustain ATP production on the timescale of synaptic plasticity. Resolving this required combining a correlative light and electron microscopy pipeline with deep-learning segmentation, DNA-PAINT, and ATP imaging in single spines and mitochondria. Following synaptic plasticity, juxtaspinal mitochondrial regions enlarge, and their local ATP output rises. Within these regions, cristae become more complex, with greater cristae membrane area, more crista junctions, and higher curvature, alongside more MIC60 and ATP synthase, increased ER contacts, and more mitochondria-proximal ribosomes. Because dendritic mitochondria are long, this remodeling occurs within one region of a mitochondrion rather than across the whole organelle, a spine-specific, sub-organellar specialization, requiring matrix-PKA activity, that expands metabolic capacity locally to fuel plasticity. More broadly, because local mitochondrial volume tracks spine-cluster size even at the modest resolution of connectomics datasets (Fig. 1H-J, S1M), these structural signatures may serve as readouts of local energy demand and of a neuron’s recent activity or plasticity history.

Existing deep-learning tools for electron tomography have been developed largely for cryo-ET for single structures, and to our knowledge none has jointly segmented multiple mitochondrial subregions with the surrounding ER and ribosomes in plastic-section tomograms while enforcing their biological relationships^54-56^. Our model (Fig. S4) generated all five labels in ∼100 s per tomogram and ∼70 min for 42 tomograms on a single workstation rather than dedicated GPU infrastructure, although differences in modality, hardware, and proofreading between studies make these time comparisons contextual rather than a normalized benchmark. Because neither our automated spine-base definition nor the segmentation framework is specific to mitochondria, both can be applied to other organelles, cell types, and disease models, and retrained for additional structures.

How is mitochondrial remodeling coupled to demand? Soluble adenylyl cyclase (sAC) produces matrix cAMP in response to metabolic state^47,49^, positioning intramitochondrial cAMP-PKA to sense energy status and drive remodeling. Because sAC is activated by bicarbonate and Ca^2+^, whether this remodeling is calcium-dependent is a key question. In single-spine plasticity, acute local mitochondrial ATP synthesis is driven by activity-evoked mitochondrial calcium uptake^3^, whereas homeostatic plasticity is induced by silencing input activity, with a fall in postsynaptic calcium influx serving as the trigger^15,57^. Juxtaspinal mitochondrial remodeling therefore occurs under opposite calcium conditions in the two paradigms, implying that cAMP-PKA-driven remodeling is either calcium-independent (for instance, by sensing metabolic state through bicarbonate) or occurs over slower timescales than the fast, calcium-triggered ATP response.

Our timing data indicate that juxtaspinal mitochondrial remodeling develops more slowly than the ATP response. During single-spine plasticity, juxtaspinal ATP rises within 2 minutes, before mitochondrial enlargement becomes statistically detectable at 17 minutes. The immediate calcium-driven rise is anticipatory^3^, arriving ahead of both the demand it will meet and the structural change that follows; remodeling is therefore positioned to support sustained output rather than to initiate it. Over 24 hours of homeostatic plasticity, the two are not temporally separable, so whether this sequence is general or specific to acute, single-spine plasticity remains open.

We further find that matrix PKA is required for MIC60 phosphorylation. This phosphorylation is a plausible integration point for coupling demand to remodeling, but its consequences are context-dependent: PINK1 phosphorylation of MIC60 drives its oligomerization and crista junction formation^7^, whereas PKA phosphorylation of MIC60 at Ser528 can instead destabilize MICOS assembly during mitochondrial damage^45^. Because we detect MIC60 phosphorylation as a PKA-dependent Phos-tag mobility shift rather than with a site-specific antibody, the modified residue(s) remain unmapped; whether this reflects Ser528 or another residue is unknown, though the matrix PKA dependence points to MIC60’s short matrix-exposed N-terminus rather than its IMS-facing coiled-coil and mitofilin domains^58,59^. Mapping this site and testing parallel shapers such as starvation-activated OPA1 oligomerization^60^, are important next steps.

Several observations support the specificity of our PKA perturbation. PKI is PKA-selective but not specific: at high concentrations it increases, rather than inhibits, the activity of several kinases, including PKC isoforms, ROCK1, and p70S6K^50,61^. These off-target effects require micromolar PKI, more than three orders of magnitude above the sub-nanomolar concentrations that inhibit PKA; and increased kinase activity would not reproduce our loss-of-mitochondrial-enlargement phenotype (Fig. 6D, F). Furthermore, Matrix-PKI is compartment-restricted, leaving cytosolic PKA substrates unaffected (Fig. S12A-D). PKA can also phosphorylate electron-transport chain subunits in non-neuronal cells and isolated mitochondrial systems, where PKA inhibition reduced baseline cytochrome c oxidase activity^47,49^. Whereas Matrix-PKI leaves baseline ATP and resting mitochondrial area unchanged (Fig. 1F, 2D, 6D, E). Matrix-PKI also blocks juxtaspinal mitochondrial enlargement (Fig. 6D, F), which a direct effect on the electron transport chain cannot explain. The loss of remodeling therefore provides the most parsimonious account of the reduced ATP and plasticity, though fully separating the structural and bioenergetic contributions of MIC60 phosphorylation will require a site-specific phosphomutant.

High-curvature cristae concentrate ATP synthase^26^. The increased curvature and ATP synthase number we observe, together with the rise in mitochondrial ATP, support expanded bioenergetic capacity during plasticity. In other systems, ATP synthase dimers and their arrays generate this curvature^28,62^. Although we had aimed to resolve these dimers and arrays, immunolabeling efficiency and DNA-PAINT acquisition did not achieve the required resolution because the resolution of our 3D DNA-PAINT (∼13 nm lateral; ∼60 nm axial) approaches the dimer spacing (12-13 nm^63,64^), and antibody labeling is subject to steric hindrance in tightly packed assemblies; therefore, not all molecules may be detected. The localizations we detect, therefore, miss higher-order assemblies that cryo-electron tomography could resolve directly.

The coordinated increase in MERCs and mitochondria-proximal ribosomes fits growing evidence that MERCs supply lipids for membrane biogenesis and that local protein synthesis maintains and remodels organelles^2,5,65-67^. Because our measurements were made 50-150 μm from the soma, they raise the possibility that ribosomes supply proteins for mitochondrial remodeling near distal spines^14^.

Overall, our findings suggest that the mitochondrion is not a uniform power supply but a mosaic of sub-compartments that adapt independently, expanding their energy-producing machinery precisely where and when demand arises. Energy allocation within a neuron is therefore regulated at a finer spatial scale than the organelle itself, a logic that may extend to other organelles and to any cell with similarly polarized architecture. Synaptic plasticity is organized in clusters along dendritic segments^68,69^, which requires a local gate that sets which synapses can change, and over what distance. Mitochondria are positioned to serve as that gate, and our measurements suggest it is built from nested spatial scales. Mitochondrial structural remodeling is confined to the juxtaspinal region (2 μm), the ATP it generates spreads ∼10 μm to neighboring spines^3^; and the stable mitochondrial compartments span ∼30 μm, matching the segment over which clustered plasticity occurs, and whose disruption blocks plasticity in neighboring spines^1,2,68,70^. The source is therefore spine-specific, while the supply serves a cluster, allowing remodeling at one spine to license plasticity across its neighbors. Conversely, failure of local energy allocation, rather than a global energy deficit, may underlie the synaptic vulnerability that accompanies mitochondrial ultrastructural defects in aging and neurodegeneration.

## Acknowledgments

We thank C. Hanus for the PSD95-mCherry plasmid; J. de Juan-Sanz for the Homer2-mOrange2-luc plasmid; G. Ashrafi for the mito-pHluorin plasmid; and H. Bito for the RCaMP1.07 plasmid. PSD95FingR-EGFP plasmid was a gift from D. Arnold (Addgene 46295). We are thankful to S. Perez, D. Bhuyan, J. Patarroyo, O. Bapat, and R. Fan for preparing cultured hippocampal neurons and for technical assistance; D. Guerrero-Given and C. I. Thomas at the MPFI Imaging Center Core for help with the transmission electron microscope; C. Bouchet-Marquis from ThermoFisher for the initial dual-axis tilt series acquisitions; N. Rzepka (scalable minds) for assistance with Webknossos and fast accessibility of the somatosensory connectomics dataset; J. Yu at the MPFI Molecular Virology Core for assistance with molecular cloning; J. Kuhl for the illustration, and B. D. Mensh, H. Inagaki, K. Harris, D. Fitzpatrick, and all members of the Rangaraju Lab for critical input on the manuscript. CS is supported by the Lundbeck Foundation (Lundbeckfonden grant no. R361-2020-2654), the Novo Nordisk Foundation (NNF23OC0085864), the Danish National Research Foundation (DNRF133), and Independent Research Fund Denmark (Danmarks Frie Forskningsfond; 4283-00041B and 5253-00010B). IG is funded by the Carl Angus DeSantis Foundation; VR is supported by the Max Planck Society, Louis D. Srybnik Foundation, F.O.R.E Foundation, Chan Zuckerberg Initiative DAF an advised fund of the Silicon Valley Community Foundation grant numbers 2023-331775 and 2024-349543, and the NIH Director’s New Innovator Award (DP2 MH140148-01).

## Author contributions

VR and MS designed experiments. Experiments were carried out by MS, IG, NSR, and TP. Data were analyzed by MS, IG, NSR, and LP. IG performed the single-spine plasticity induction, ATP and pH imaging experiments, and analysis. LP performed the early MICrONS connectomics data analysis, while SL supervised the mouse somatosensory connectomics data analysis. VV performed the preliminary DNA-PAINT experiments that are not included in the manuscript, and CS supervised the DNA-PAINT experiments and analysis. TP performed the electrophysiology experiments, and RY supervised them. NK supervised the electron microscopy sample preparation and imaging. All other experiments were conducted and analyzed by MS. VR supervised all experiments and analysis. VR and MS wrote the manuscript, and all authors reviewed and edited it.

## Methods

### Animals

All experiments were performed in accordance with the Max Planck Florida Institute for Neuroscience IACUC regulations (protocol number 25-004).

### Plasmid constructs

The PSD95FingR-mTagBFP2 plasmid was made by replacing EGFP with mTagBFP2 in the PSD95FingR-EGFP plasmid^21^, purchased from Addgene (Plasmid no. 46295). The OMM-EGFP plasmid was made by replacing GCaMP6f in the OMM-GCaMP6f plasmid^71^. The MPC1 shRNA (Plasmid no. 229016), and GCaMP6s (Plasmid no. 40753) were purchased from Addgene. For the Matrix-PKI and IMS-PKI constructs: the PKI sequence was taken from pRSV-PKI-V2 (Addgene plasmid #45066) and added to the backbone of Mito-BFP (Addgene plasmid #49151) for Matrix-PKI; and the pre-CoxIV sequence in Matrix-PKI was replaced with the LactB sequence from IMS-APEX2 (Addgene plasmid #79058) for IMS-PKI.

### Cell culture preparation and transfection

Unless specified otherwise, all reagents were purchased from Sigma, and all stock solutions were stored at -20 °C. Timed-pregnant Sprague-Dawley rats were obtained from Charles River Laboratories and housed at the animal core facility of the Max Planck Florida Institute for Neuroscience to obtain the postnatal P0 pups subsequently. Hippocampal regions were dissected in ACSF containing (in mM) 124 NaCl, 5 KCl, 1.3 MgSO4:7H2O, 1.25 NaH2PO4:H2O, 2 CaCl2, 26 NaHCO3, and 11 Glucose (stored at 4 °C) and stored in hibernate E buffer (BrainBits LLC, stored at 4°C). Dissected hippocampi were dissociated using the Papain Dissociation System (Worthington Biochemical Corporation, stored at 4 °C) with a modified manufacturer’s protocol. Briefly, hippocampi were digested in papain solution (20 units of papain per ml in 1 mM L-cysteine with 0.5 mM EDTA) supplemented with DNase I (final concentration 95 units per ml) and placed in a shaking heat block for 30 min at 37 °C, 900 rpm. Digested tissue was triturated and set for 5 min, and the supernatant devoid of tissue chunks was collected. The supernatant was centrifuged at 300 g (0.3 rcf) for 5 min, and the pellet was resuspended in resuspension buffer (1 mg of ovomucoid inhibitor, 1 mg of albumin, and 95 units of DNase I per ml in EBSS). The cells were forced to pass through a discontinuous density gradient formed by the resuspension buffer and the Ovomucoid protease inhibitor (10 mg per ml) with bovine serum albumin (10 mg per ml) by centrifuging at 70 g (0.6 rpm) for 10 min at room temperature. The final cell pellet devoid of membrane fragments was resuspended in Neurobasal-A medium (Gibco, stored at 4°C) supplemented with Glutamax (Gibco, stored at -20 °C) and B27 supplement (Gibco, stored at -20°C) and 3% fetal bovine serum (R&D Systems, stored at -20 °C). For ATP imaging experiments, we used a serum-free plating medium to obtain a detectable basal ATP signal for luminescence imaging. Cells were plated on poly-D-lysine-coated coverslips mounted on MatTek dishes at a density of 70000-80000 cells/cm^2^. All coverslips were 35 mm in diameter and 1.5 mm thick, with alphanumeric labels for the CLEM experiments. For DNA-PAINT experiments, high-precision MatTek dishes with a coverslip diameter of 35 mm and a thickness of 0.17 mm were used. Cultures were maintained at 37 °C and 5% CO2 by feeding them with the same medium every 3 - 4 days until transfection. Transfections were performed 12 -18 days after plating by magnetofection using Combimag (OZ biosciences, stored at 4°C) and Lipofectamine 2000 (Invitrogen, stored at 4°C) according to the manufacturer’s instructions.

### Imaging and optical system

Live cell imaging was conducted between 17-21 days after plating. All experiments were performed at 37 °C in modified E4 imaging buffer containing (in mM) 120 NaCl, 3 KCl, 10 HEPES (buffered to pH 7.4), 2 CaCl_2_ (4 CaCl_2_ for single-spine plasticity induction with 2-photon glutamate uncaging), 0.81 MgCl_2_ (no MgCl_2_ for single-spine plasticity induction with 2-photon glutamate uncaging) and 10 Glucose, in the absence of TTX, even for homeostatic scaling-induced neurons, unless specified otherwise.

Imaging was performed using a custom-built inverted spinning disk confocal microscope (3i imaging systems; model CSU-W1) with two cameras: an Andor iXon Life 888 electron multiplying charge-coupled device (EMCCD) camera for confocal fluorescence imaging and an Andor iXon3 888 camera for luminescence imaging. The Andor iXon3 888 EMCCD camera was selected for ultralow dark noise, further reduced by cooling to −100 °C. The speed of the Andor iXon3 888 camera used for luminescence measurements was 1 MHz with 1.00 gain, 1 vertical speed, and 1000 intensification. Image acquisition was controlled by the SlideBook 6 software. Images were acquired with a Plan-Apochromat 63X/1.4 NA. Oil objective, M27 with DIC III prism, using a CSU-W1 Dichroic for 488/561 nm excitation with Quad emitter and individual emitters, at laser powers 50 ms exposure, 4.3 mW (488 nm) and 100 ms exposure, 5.5 mW (561 nm) for confocal fluorescence, and a 720 nm multiphoton short-pass emission filter for luminescence. During imaging, the temperature was maintained at 37 °C using an Okolab stage top incubator with temperature control.

### Single-spine plasticity induction, imaging and analysis

For single-spine plasticity induction experiments, rat hippocampal neurons expressed Mito-EGFP and RCaMP1.07, with Matrix-PKI-mTagBFP2 where specified. RCaMP1.07 fluorescence was used to identify dendrites and spines and monitor spine-specific calcium responses, while Mito-EGFP was used to visualize mitochondria. Matrix-PKI-expressing neurons were identified from the mTagBFP2 signal. Before two-photon glutamate uncaging and imaging, the imaging buffer was replaced with modified E4 buffer lacking Mg²⁺ and containing 2 μM TTX (citrate salt; 2 mM stock prepared in water), 50 μM forskolin (Tocris Bioscience; 25 mM stock prepared in DMSO), and 2 mM 4-methoxy-7-nitroindolinyl-caged-L-glutamate (MNI-caged glutamate; Tocris Bioscience; 100 mM stock prepared in E4 buffer).

Two-photon glutamate uncaging was performed using a Mai Tai HP multiphoton laser at 720 nm. Uncaging was performed at 7.5 mW laser power measured at the back aperture. Spines located at least 50 μm from the cell body were selected. Because neurons were derived from dissociated cultures, apical and basal dendrites were not distinguished. Spines positioned near a single, unbranched mitochondrial compartment were selected independently of spine morphology. To verify spine responsiveness, an uncaging spot was positioned adjacent to the spine head and 1–2 test uncaging pulses (10 ms per pixel) were delivered. Spines exhibiting a detectable calcium response were used for subsequent plasticity induction. Synaptic plasticity was induced using 60 uncaging pulses delivered at 0.5 Hz, with a 10-ms pulse duration per pixel and 7.5 mW laser power.

For structural measurements, 6-μm Z-stacks were acquired with a 0.1-μm step size at 1024 × 1024 pixels and a lateral sampling of 0.2 μm per pixel. Baseline stacks were acquired at 1-min intervals at t = −2, −1, and 0 min. Plasticity was then induced over 2 min, followed by an immediate post-stimulation acquisition at t = 2 min and subsequent acquisitions every 5 min for up to 62 min. Three-dimensional stacks were maximum-intensity projected and the resulting time series were registered to correct XY displacement between time points. Before segmentation, maximum-projected images were upsampled in XY to provide a finer spatial sampling grid for boundary placement. This interpolation did not increase the underlying optical resolution, but reduced quantization of segmentation boundaries to the native 0.2-μm pixel grid and enabled subpixel boundary placement for area measurements, analogous to manually drawing object boundaries across image pixels in Fiji while avoiding observer-dependent boundary selection.

Mitochondria were segmented from the upsampled maximum-projected Mito-EGFP images using a custom deep-learning model. Upsampling did not increase optical resolution, but provided a finer sampling grid on which segmentation boundaries could be represented, reducing pixel-edge quantization in area measurements and permitting subpixel boundary placement without the observer-dependent variability associated with manual tracingFor each stimulated spine and time point, a line crossing the dendrite perpendicular to its long axis was manually marked in Fiji at the spine base when clearly visible, or directly beneath the spine head when the base could not be unambiguously visualized. This line provided a local reference for orientation of a fixed 2 × 4 μm juxtaspinal region, with the 2-μm dimension oriented along the dendritic axis. Mitochondrial area was calculated as the area of the Mito-EGFP segmentation contained within this fixed region, providing a standardized measurement of juxtaspinal mitochondrial area across time. Where non-juxtaspinal regions were analyzed, equivalent fixed regions were positioned >2 μm away from the juxtaspinal region within the same mitochondrial compartment. Juxtaspinal mitochondria were also manually categorized according to the morphological changes observed following plasticity induction as recruitment, protrusion, budding, or no detectable change.

Spine-head width was quantified from RCaMP1.07 fluorescence using the same image series. At each time point, a line was manually drawn across the spine head in Fiji and the fluorescence-intensity profile along the line was fitted with a Gaussian function. Spine-head width was defined as the full width at half maximum (FWHM) of the fitted Gaussian. This intensity-profile measurement allowed changes in spine-head width to be quantified independently of absolute RCaMP1.07 fluorescence intensity.

### Mitochondria and spine imaging and analysis following homeostatic plasticity

For homeostatic plasticity experiments, mitochondria were labeled with Mito-EGFP and postsynaptic puncta were labeled with PSD95-mCherry or PSD95FingR-EGFP, as specified. Homeostatic plasticity was induced with 2 μM TTX. Measurements were performed either in independent biological replicates imaged after 24 h of Control or TTX treatment, or longitudinally over 24 h in the same neurons. For longitudinal imaging, a baseline Z-stack was acquired before TTX addition (t = −0.25 h), followed by acquisition immediately after TTX addition (t = 0 h) and subsequently every hour for 24 h. Control neurons were imaged using the same acquisition schedule in the absence of TTX.

For mitochondrial and spine structural analysis, 6-μm Z-stacks were acquired with a 0.1-μm axial step size at 1024 × 1024 pixels and a lateral sampling of 0.2 μm per pixel. For time-series experiments, stacks from each time point were maximum-intensity projected and registered in XY before further analysis. Single-time-point stacks were similarly maximum-intensity projected. The maximum projections were then upsampled in XY before segmentation. The resulting images were supplied to separate custom deep-learning models trained to segment Mito-EGFP-labeled mitochondria and PSD95-labeled postsynaptic puncta, respectively.

To restrict measurements to the dendritic compartment, dendritic arbors were manually traced from the PSD95 channel in Fiji using the segmented line tool. The resulting arbor geometry was used as a spatial reference, and only segmented mitochondria and PSD95 puncta located within ±6 μm of the dendritic arbor were retained for subsequent analysis. This excluded structures spatially separated from the analyzed dendrites while preserving mitochondria and spines associated with the local dendritic segment.

For 24-h endpoint experiments, PSD95 puncta that remained detectable throughout the 24-h time series were randomly selected manually in the first frame. The selected puncta were subsequently tracked between frames from the PSD95 segmentation using a linear-assignment-problem (LAP) tracker, thereby maintaining the identity and position of individual spines throughout the experiment.

Juxtaspinal mitochondrial area was measured within a fixed 2 × 4 μm region positioned at the base of each selected spine. The local dendritic arbor geometry was used to automatically orient the region such that the 2-μm dimension followed the dendritic axis. Mitochondrial area was calculated from the portion of the deep-learning-derived Mito-EGFP segmentation contained within this fixed region. Use of an identically sized, dendritically oriented sampling region allowed mitochondrial area to be compared between spines, conditions, and time points. For non-juxtaspinal measurements in the 24-h endpoint experiments, all segmented PSD95 puncta were mapped to the corresponding dendritic arbor. Dendritic regions containing at least 2 μm of arbor without associated PSD95 puncta were identified as non-juxtaspinal regions, and the same 2 × 4 μm sampling region was positioned and oriented using the local dendritic geometry. Mitochondrial area within the Mito-EGFP segmentation was then measured identically for juxtaspinal and non-juxtaspinal regions.

Spine area was measured directly from the PSD95 segmentation for all spines. In longitudinal experiments, the tracked puncta identity was used to measure the same spine across successive time points, allowing measurement of juxtaspinal mitochondrial area over the 24-h period.

### Deep learning-based segmentation of mitochondria and PSD95 labeled spines in fluorescence images

We developed separate deep-learning models to segment Mito-EGFP-labeled mitochondria and PSD95-labeled postsynaptic puncta from the maximum-projected fluorescence images. Ground-truth masks were generated by manual segmentation in ORS Dragonfly. Before model input, images were upsampled in XY to provide a finer sampling grid for object boundaries. This interpolation was not meant to increase optical resolution but reduce quantization of segmentation boundaries to the native pixel grid and enabled subpixel boundary placement without manual tracing.

Three features were generated from each frame: globally z-score-normalized fluorescence, locally normalized fluorescence calculated from a Gaussian-filtered local mean and variance (σ = 12 pixels), and a boundary-sensitive feature obtained from the two-dimensional gradient magnitude of the locally normalized image after Gaussian smoothing (σ = 1 pixel). Normalized intensities were limited to ±6 standard deviations. For time-series datasets, four additional series-level features were generated by averaging the global normalization, local normalization, and gradient features across time and by calculating the fraction of frames in which locally normalized fluorescence exceeded 0.5 at each pixel.

The model consisted of two coupled two-dimensional residual U-Nets. A series-level network used the four time-series features to generate a spatial guidance probability map, which was supplied to a second network together with the three features from each individual frame. The guidance map was used both as an additional input and to spatially modulate decoder skip features, allowing structures consistently present throughout the time series to guide segmentation while preserving frame-specific boundary changes. Both networks contained four encoder–decoder resolution levels with residual blocks comprising two 3 × 3 convolutions, group normalization, and SiLU activation. Residual projections used 1 × 1 convolutions when channel dimensions differed. Downsampling used 2 × 2 max pooling and upsampling used transposed convolutions. The guidance and frame-specific networks used 16 and 24 initial feature channels, respectively.

Training data were divided into 80% training and 20% validation sets using a fixed random seed. Training augmentation included horizontal and vertical reflections, 90° rotations, fluorescence scaling and offsets, gamma transformations, and Gaussian noise. Foreground pixels were weighted in the binary cross-entropy loss according to the background-to-foreground ratio, capped at 50. The frame-specific loss was the sum of weighted binary cross-entropy and Dice losses. The guidance branch was trained against a soft occupancy target generated from the mean ground-truth mask across the image series, Gaussian-smoothed with σ = 3 pixels, using binary cross-entropy and soft-Dice losses; this loss was weighted by 0.2 relative to the frame-specific loss. Models were trained in PyTorch using AdamW (learning rate, 1 × 10⁻⁴; weight decay, 1 × 10⁻⁵; batch size, 1), and the checkpoint with the lowest validation loss was retained. Probability maps were binarized at a threshold of 0.5.

Segmentation performance was evaluated using Dice coefficient, intersection-over-union (IoU), precision, recall, and pixel-wise accuracy, with Dice and IoU considered the primary metrics because of the large foreground–background imbalance. The retained Mito-EGFP model achieved a Dice coefficient of 0.860, IoU of 0.754, precision of 0.786, recall of 0.950, and accuracy of 0.998. The retained PSD95 model achieved a Dice coefficient of 0.840, IoU of 0.724, precision of 0.795, recall of 0.890, and accuracy of 0.999.

### Electrophysiology measurements and analysis

Rat hippocampal neurons cultured on 12-mm glass coverslips were transferred to the stage of an Olympus BX51WI microscope and maintained at room temperature in a recording buffer (identical to the E4 imaging buffer, osmolarity: 305–310 mOsm). Neurons expressing Mito-EGFP (mitochondria) and PSD95-mCh (spines), along with Matrix-PKI where specified, were identified via epifluorescence using a 470 nm and 561 nm LED light source and an X-Cite® 120Q illumination system (Excelitas, PA). In experiments containing Matrix-PKI, the BFP signal in the Matrix-PKI construct could not be directly assessed because the electrophysiology imaging system lacked a 405 nm excitation channel. Therefore, Mito-EGFP-expressing neurons in the Matrix-PKI experimental group were assumed to also express Matrix-PKI.

Whole-cell voltage-clamp recordings were performed using patch pipettes (3–5 MΩ) pulled from borosilicate glass (Sutter Instruments) and filled with an internal solution containing (in mM): 145 K-gluconate, 14 phosphocreatine, 4 NaCl, 0.3 NaGTP, 4 MgATP, 3 L-ascorbic acid, and 10 HEPES (pH 7.3, 294 mOsm). Seal resistances ranged from 2–6 GΩ. Recordings were excluded if the series resistance exceeded 25 MΩ; series resistance was left uncompensated.

After establishing whole-cell configuration, neurons were held at −70 mV, and 3– 5-minute-long recordings were acquired beginning 3–5 minutes post-break-in. Data were recorded using a MultiClamp 700B amplifier (Molecular Devices) and digitized at 10 kHz with custom C# software (FLIMage; https://github.com/ryoheiyasuda/FLIMage_public) on a Windows PC. Neurons with resting membrane potentials more positive than −50 mV were excluded from analysis.

Recordings were low-pass filtered at 1 kHz using a Bessel filter in ClampFit (version 11.4.2.04, Molecular Devices). mEPSC events were detected using the template search function, and only events with amplitudes ≥ 5 pA were included. All mEPSC events from each neuron were pooled by condition (e.g., control or homeostatic plasticity) and analyzed as cumulative distributions representative of the condition.

### Local mitochondrial volume quantification from somatosensory L4 and visual cortex L2/3 connectomics datasets

Three-dimensional reconstructions of neuronal and mitochondrial meshes were obtained from the MICrONS Phase 1 mouse visual cortex layer 2/3 dataset, together with annotation tables containing mitochondrial IDs, neuronal IDs, synapse IDs, synaptic cleft size (cleft_vx; voxel dimensions, 4 × 4 × 40 nm), and synapse coordinates^20^. For the mouse somatosensory cortex layer 4 dataset^18^, with an effective segmentation resolution of approximately 11.24 × 11.24 × 28 nm, neuronal and mitochondrial surface meshes were generated from the corresponding volumetric segmentations using marching cubes^72,73^. Synapse IDs and agglomeration IDs were obtained from the accompanying annotation tables, with each agglomeration ID treated as the neuronal identifier analogous to the neuronal IDs used in MICrONS.

To quantify juxtaspinal mitochondrial volume within connectomics datasets, we developed a custom pipeline for accurate estimation of the spine base (i.e., the point from which a spine originates from the dendritic shaft). Automated estimation of the spine base is a nontrivial process in complex three-dimensional reconstructions because skeleton branch points do not necessarily coincide with the anatomical transition between the narrow spine neck and the broader dendritic shaft. Classical automated approaches have instead identified this boundary from changes in voxel spread or by segmenting complete spine volumes^74,75^, while recent connectomics frameworks use mesh or point-cloud segmentation to classify spine heads, necks, and shafts^76,77^. We developed a geometry-adaptive approach that determines the spine base directly from the caliber transition along a synapse-anchored centerline, avoiding the requirement to separately segment each spine into head, neck, and dendritic compartments.

For each postsynaptic site, the synaptic coordinate was mapped to the neuronal surface and a 7-µm geodesic neighborhood was extracted. The local mesh was skeletonized from the synaptic position using exact wavefront skeletonization in Skeletor with a 30-nm step size. Wavefront skeletonization collapses successive geodesic level sets into a centered curve representation and provides a local radius at each skeleton node^78^. To identify the local dendritic shaft, skeleton nodes with radii <100 nm were excluded and the longest path through the remaining skeleton was retained. The shortest graph trajectory connecting the synaptic position to this dendritic path then provided a one-dimensional representation of the spine-to-shaft transition.

An initial spine-base estimate was placed along this trajectory at a distance from the dendritic meeting point equal to the local dendritic radius. We then refined the base from the increase in skeleton radius that occurs as the relatively narrow spine neck enters the wider dendritic shaft. Beginning near the initial estimate and proceeding toward the dendrite, the radius profile across ordered skeleton nodes was fitted with a rectified-linear function,

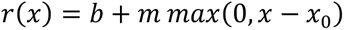

where (*b*) represents the local neck-radius baseline, (*m* ≥ 0) the increase in radius toward the dendrite, and (*x*_0_) the onset of this increase. The observed skeleton node nearest the fitted breakpoint (*x*_0_) was designated as the final spine base. When fewer than four nodes were available for fitting, the dendritic-radius estimate was retained. Thus, rather than defining the spine base from a raw skeleton bifurcation or a fixed distance threshold, the algorithm identifies the local geometric transition between spine and shaft and therefore adapts to differences in spine-neck and dendritic caliber.

After the spine-base coordinate had been fixed, the synapse-to-base trajectory was refined using an LST-inspired anisotropic Laplacian relaxation adapted from geometry-preserving smoothing approaches^79-82^. At each iteration, displacement of each interior node toward the mean position of its neighboring nodes was decomposed into components normal and tangential to the local trajectory. Normal displacement was retained while tangential displacement was damped, reducing local irregularities introduced by mesh triangulation and wavefront discretization without allowing substantial redistribution of nodes along the spine axis. Relaxation was performed for 100 iterations with a step size of 0.5 and tangential damping of 0.5, while the synaptic and ReLU-defined spine-base endpoints remained fixed. The resulting relaxed centerline preserved the anatomically determined endpoints while providing a smooth trajectory for subsequent distance and spatial measurements.

To enable connectome-scale analysis, spine-base estimation was performed locally and in parallel. The neuronal surface graph was constructed once for each neuron, after which independent geodesic neighborhoods and spine trajectories were calculated concurrently for multiple synapses. Smaller neuronal meshes were processed in parallel across neurons, whereas larger meshes were processed individually to limit memory use. Restricting analysis to local surface graphs avoided repeatedly processing the underlying dense image volumes and reduced each spine to a compact centerline-and-radius representation. This provided a scalable route to automatically determine spine-base coordinates across large neuronal reconstructions while retaining the local three-dimensional geometry required for subsequent mitochondrial measurements.

Mitochondrial meshes were separated into connected components and skeletonized using the same wavefront framework. For each component, the longest terminal-to-terminal path and connected branches ≥ 1 µm were retained. These mitochondrial centerlines were also subjected to LST-inspired anisotropic relaxation, with terminal and branch points constrained, before morphometric measurements were made. Each mitochondrial surface vertex was assigned to the nearest relaxed centerline node belonging to the same component, and local mitochondrial diameter was calculated as twice the mean distance between each centerline node and its associated surface vertices. Local diameters were normalized within each mitochondrial component as z-scores, thereby identifying changes in caliber relative to the intrinsic width of that mitochondrion rather than imposing a common absolute diameter threshold. Regions with normalized width >1 were classified as enlarged, and neighboring enlarged regions within 200 nm were grouped into individual remodeling domains.

Upon manual inspection of the neuronal meshes in these intact brain datasets, we observed that neighboring spines frequently sampled the same juxtaspinal mitochondrial region when their bases originated from nearby points along the dendritic shaft. Additionally, more than one synapse ID sometimes shared the same spine structure and therefore had a common spine base. To control for these anatomical variations prominently present within densely-innervated intact brain samples, we sampled a common juxtaspinal mitochondrial region for groups of spines with overlapping juxtaspinal regions. We first projected spine bases onto the dendritic centerline and clustered them using DBSCAN with a 1 μm spatial radius. The centroid of each cluster was projected back onto the dendritic centerline to provide a common juxtaspinal reference point. Total mitochondrial volume was then measured within a 2 μm-diameter sphere centered on the reference point. To estimate the total spine size associated with each juxtaspinal region, we summed the synaptic cleft segmentation volume within each spine cluster for MICrONS and the axon–spine interface (ASI) surface area for the somatosensory cortex L4 dataset. Thus, the visual cortex dataset measurement represents the volume of the segmented synaptic interface voxels, whereas the somatosensory cortex dataset measurement represents the surface area of the corresponding axon–spine interface.

The somatosensory cortex L4 reconstructions were additionally used to identify mitochondrial morphologies corresponding to those observed experimentally. A spine was considered to have a nearby mitochondrion when the nearest mitochondrial surface was within 2 μm of its base. Mitochondrial enlargement was defined as a normalized local diameter >1. Protrusion of juxtaspinal mitochondrial regions was defined by a normalized diameter >1.5 together with a spine-base-to-mitochondrion surface distance <200 nm, thereby identifying enlarged mitochondrial regions protruding toward the spine base. Because dynamic recruitment of mitochondrial content cannot be measured in static reconstructions, recruitment was calculated as enlarged cases not classified as protrusions. While this definition does not establish recruitment of mitochondrial content as observed in our live-neuron experiments, it provides a crude estimate of mitochondrial enlargement that is larger than the rest of the mitochondrion but not protruding into the spine base. Mitochondria present within the spine head, analogous to budding observed in live-neuron experiments, were identified independently of mitochondrial enlargement. A mitochondrion was classified as being within a spine when the synapse-to-mitochondrial-surface distance was within 500 nm and the mitochondrion present in the spine had a distinct volume from that present at the spine base. The frequencies of mitochondrial enlargement, protrusion, recruitment, and budding, were then quantified across the total reconstructed synapse population.

### ATP_mito_ and ATP_spine_ imaging and analysis

Rat hippocampal neurons were transfected with Mito-ATP (Mito-mCh-Luc) and PSD95FingR-EGFP, along with either MPC1 shRNA or Matrix-PKI where specified, for imaging mitochondrial ATP and spines, respectively. Neurons were transfected with Spn-ATP (Homer2-mOrg-Luc) and Mito-EGFP, along with either MPC1 shRNA or Matrix-PKI where specified, for imaging spine ATP and mitochondria, respectively. For homeostatic plasticity induction, the transfected neurons were incubated with TTX (citrate salt; 2 μM) for 24 h in culture medium. Before ATP imaging, we replaced the imaging buffer with 2 mM luciferin (Promega) prepared in imaging buffer to enable bioluminescence detection of ATP in mitochondria and spines. Transfected neurons were identified by mCh fluorescence of the Mito-ATP or mOrg fluorescence of the Spn-ATP. Test images were acquired simultaneously in the 561 nm (mOrg/mCh) and 488 nm (PSD95FingR-EGFP/Mito-EGFP) channels at exposure times of 100 ms for the 561 nm and 50 ms for the 488 nm channels, and bioluminescence images were simultaneously acquired over 60 s for Spn-ATP and 20 s for Mito-ATP. Only neurons exhibiting both mature spines and healthy dendritic mitochondria were selected for imaging. For each condition, a 3-minute 2D time series of alternating luminescence and fluorescence frames was acquired. To measure juxtaspinal and non-juxtaspinal mitochondrial ATP levels and visualize the presence of mitochondria at the base of the spines, a 3D Z-stack of the same neurons was acquired in the 561 nm and 488 nm channels, with a step size of 0.1 μm spanning a range of 6 μm, following the 2D ATP imaging.

All image analysis was performed using custom batch-processing macros in Fiji. Each time series was average-projected and background-subtracted. To correct for misalignment between confocal fluorescence and luminescence images, luminescence images were aligned to fluorescence images using the Multichannel SIFT alignment plugin.

For juxtaspinal and non-juxtaspinal Mito-ATP measurements following homeostatic plasticity induction, mitochondria were marked using composite images of the 561 nm (mCh), 488 nm (PSD95-FingR-EGFP), and luminescence (Mito-ATP) channels. Circular ROIs (2 μm in diameter) were positioned over juxtaspinal mitochondrial regions based on the presence of PSD95 puncta and a detectable, in-focus luminescence signal. Non-juxtaspinal mitochondrial ROIs were selected within the same mitochondrial compartments, using the juxtaspinal ROI as a spatial reference and positioned >2 μm away from the juxtaspinal ROI. These ROIs were confirmed to have detectable, in-focus luminescence signal and lack PSD95 puncta. Each ROI was then Otsu-thresholded to generate a mitochondrion-shaped ROI using the mCh channel at the juxtaspinal and non-juxtaspinal regions, spanning a 2 μm segment of the mitochondrion. For axonal Mito-ATP measurements in control vs. TTX-treated neurons, circular ROIs (2 μm) were positioned over axonal mitochondria with detectable, in-focus luminescence signal. The mean mCh fluorescence and corresponding luminescence intensity were then measured for each mitochondrial ROI to calculate the luminescence-to-fluorescence ratio (L/F).

To compare juxtaspinal and non-juxtaspinal ATP_mito_ following single-spine plasticity induction, we reanalyzed images from our recently published dataset^2^, in which we reported a significant increase in juxtaspinal ATP_mito_ following plasticity induction. Using the same images, we quantified ATP_mito_ in non-juxtaspinal mitochondrial regions within the same mitochondrial compartments as the plasticity-induced spines and in juxtaspinal mitochondrial regions near uninduced spines in sister dendrites of the same neurons; these measurements were not included in the original study². Circular ROIs (2 μm in diameter) were positioned over non-juxtaspinal regions, devoid of spines, within the same mitochondrial compartment, >2 μm from the juxtaspinal ROI. Juxtaspinal ROIs near uninduced spines on sister dendrites of the same neurons were similarly marked and used as control. All the circular ROIs were Otsu-thresholded, and ATP_mito_ was quantified as described above. For Spn-ATP, individual spines were manually marked in the mOrg channel using 1 μm diameter circular ROIs. The mean mOrg fluorescence and corresponding luminescence intensity were measured for each spine ROI to calculate the luminescence-to-fluorescence ratio (L/F).

### pH_mito_ and pH_spine_ measurements

Rat hippocampal neurons were transfected with Mito-pHluorin and PSD95FingR-BFP, along with either MPC1 shRNA or Matrix-PKI where specified, to image mitochondrial pH and spines, respectively. Neurons were transfected with cyto-pHluorin and Mito-DsRed, along with either MPC1 shRNA or Matrix-PKI where specified, for imaging spine pH and mitochondria, respectively. Transfected neurons were identified by pHluorin fluorescence in the 488 nm channel. Only neurons exhibiting both mature spines and healthy dendritic mitochondria were selected for imaging.

Time-lapse pHluorin imaging was performed at 3 frames per second. After ∼50 frames of imaging, the E4 imaging buffer was replaced with an NH₄Cl solution (in mM): 50 NH₄Cl, 70 NaCl, 3 KCl, 2 CaCl₂, 0.81 MgCl₂, 10 Glucose, 10 HEPES; pH adjusted to 7.4 at 37 °C.

Time series frames were aligned using the Multichannel SIFT alignment plugin in Fiji, background-subtracted, and averaged projected. For mitochondrial pH measurements, dendritic and axonal mitochondria were marked in the pHluorin channel using 2 μm diameter circles. Each circle was Otsu-thresholded to define a mitochondrion-shaped region of interest spanning a 2 μm mitochondrial segment. Mean pHluorin intensity was measured across frames for each mitochondrial ROI.

For spine pH measurements, individual spines were manually marked in the pHluorin channel using 1 μm diameter circles. Mean pHluorin intensity was measured across frames for each spine ROI.

The mean intensity of the first ten frames was used as the steady-state fluorescence intensity corresponding to the steady-state pH. The peak intensity upon NH₄Cl application was defined as the fluorescence corresponding to a pH of 7.4. The steady-state spine or mitochondrial pH values were calculated using a modified Henderson-Hasselbalch equation^3,22^ as follows:

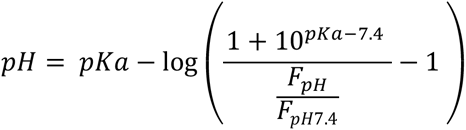

where, pKa = 7.1 for pHluorin, F_pH_ is the fluorescence intensity of pHluorin at the measured pH, and FpH_7.4_ is the peak fluorescence intensity of pHluorin at pH 7.4 upon adding NH_4_Cl.

### pH correction for ATP_spine_ and ATP_mito_ measurements

Since the pH of the spines following homeostatic plasticity was lower, pH 6.75, than the pH of the control spines, pH 6.95, the luminescence and fluorescence intensities for homeostatic plasticity were pH-corrected as follows:

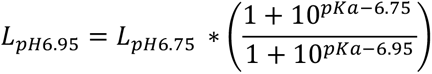

where 6.75 is the baseline pH of spines upon homeostatic plasticity, L_pH6.75_ is the luciferase luminescence intensity at pH 6.75, and pKa is 7.03 for luciferase.

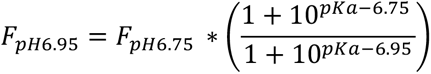

where 6.75 is the baseline pH of spines upon homeostatic plasticity, F_pH6.75_ is the mOrg fluorescence intensity at pH 6.75, and pKa is 6.5 for mOrg. L_pH6.95_/F_pH6.95_ was calculated to obtain the L/F ratio proportional to ATP_spine_. Cytosolic pH did not differ significantly between MPC1 and MPC1+TTX; therefore, we did not apply any pH correction for this dataset.

Similarly, since the mitochondrial pH did not differ significantly between control and homeostatic plasticity, and between MPC1 and MPC1+TTX, we did not pH-correct for this dataset for ATP_mito_.

However, the mitochondrial pH of Matrix-PKI+TTX (pH 6.98) was significantly less than the Matrix-PKI control (pH 7.05). Therefore, the L/F value corresponding to ATP_mito_ for PKI+TTX was pH-corrected with respect to the mito pH of the Matrix-PKI control as follows:

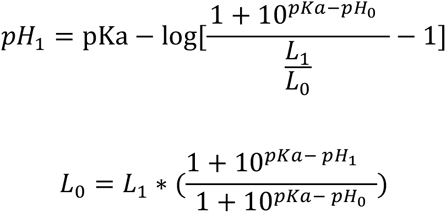

where pKa is 7.03 for ATP_mito_, pH_0_ is the average pH for PKI control, pH_1_ is the pH following homeostatic plasticity, L_0_ is the luminescence at pH_0_, and L_1_ is the luminescence at pH_1_. We used the average pH for each condition for correction, and we did not propagate errors (∼2%) from pH measurements into the final error for L/F determination.

### Electron microscopy sample preparation

Neuronal cultures were fixed following live-cell imaging for 2 hours at room temperature in a 2% paraformaldehyde (Electron Microscopy Sciences) and 2.5% glutaraldehyde (Electron Microscopy Sciences) solution, prepared in 0.14 M or 0.17 M cacodylate buffer (pH 7.4 and concentration adjusted to match the osmolarity of the E4 imaging buffer). After fixation, the dishes were washed in cacodylate buffer for 2 hours at room temperature. Samples were post-fixed by incubation with 1 ml of 2% osmium tetroxide (OsO₄) (Electron Microscopy Sciences) in cacodylate buffer for 1 hour on ice. After the initial incubation, 1 ml of 5% potassium ferrocyanide (Sigma Aldrich) was added directly to the OsO₄ solution in the dish, yielding a final working concentration of 1% OsO₄ and 2.5% potassium ferrocyanide. Neuronal samples were incubated with this mixture for an additional 1 hour on ice. The solution was then removed and replaced with 1 ml of fresh 2% OsO₄, and samples were incubated for another 1 hour on ice. Following fixation, neuronal samples were washed three times in ultrapure water (Sigma Aldrich) for 1 hour at room temperature. Neuronal samples were then stained with 1% aqueous uranyl acetate (Electron Microscopy Sciences) for 1 hour at room temperature, followed by a 1-hour wash in ultrapure water. Neuronal samples were dehydrated through a graded ethanol series (30%, 50%, 70%, 90%, 99.5%, and 100%), with each step carried out for 30 min at room temperature with gentle shaking (30 rpm). Neuronal samples were infiltrated with Durcupan ACM resin (Sigma Aldrich) without the accelerator component C, through sequential ethanol: resin mixtures (1:3 overnight, 1:1 for 6 hours, and 3:1 overnight), followed by incubation in 100% Durcupan overnight with the accelerator component C under vacuum in a desiccator. To flatten the resin surface, a small piece of ACLAR (Electron Microscopy Sciences) film was placed on top of each well. Dishes were polymerized at 60°C for 72 hours. After polymerization, dishes were left at room temperature overnight. Orientation markings made on the coverslips during live-cell imaging were copied onto the ACLAR sheet. Coverslips were carefully removed using a feather blade (Electron Microscopy Sciences) and the resin disk (in the shape of the MatTek well) containing the embedded neuronal samples was pushed out and separated from the dish along with the marked ACLAR sheet.

### Ultrathin sectioning of the target neuron

Alphanumeric labels from the coverslip were imprinted onto the surface of the resin disk together with the cultured neurons. X and Y coordinates from live neuronal confocal imaging were extracted to generate a map of relative neuron positions. ACLAR film markings were used to orient the disk, and neurons of interest were located under a stereomicroscope. For each identified neuron, the corresponding alphanumeric position was recorded onto the map to design a cutting strategy. Boxes were drawn around neurons of interest, allowing sufficient margins to account for extraction variability. Each boxed region was carefully excised from the resin disk using a razor blade. Excised resin pieces were mounted onto separately prepared cylindrical resin blocks using cyanoacrylate (Gorilla Glue). For each neuron of interest, a trapezoidal label was placed on the confocal image to define the target region. A matching trapezoid was scratched around the neuron on the resin block and trimmed precisely using a Diatome trimming knife. Serial sections were cut at a thickness of 120 or 150 nm using a Diatome Ultra knife. Sections were collected onto formvar- or pioloform-coated (Electron Microscopy Sciences, Ted Pella, Inc.) 3 mm copper grids (Electron Microscopy Sciences) with 2 mm apertures.

### Electron tomography of target mitochondria

Transmission Electron Microscopy (TEM) correlation was performed on an FEI Tecnai G2 BioTwin. All serial sections for a target region were checked, and dendrites from a neuron of interest were visually identified using the live neuron DIC images as correlative guides. PSD95-positive spines and their corresponding juxtaspinal mitochondrial regions were identified using 63X live neuron confocal images. TEM reference images were taken at 120 kV using a Veleta CCD camera. Given that mitochondria frequently span multiple serial sections and are often absent from the section containing the spine, the section displaying the most prominent mitochondrial cross-section was selected and imaged at a ∼2 × 2 μm field of view and ∼0.48 nm pixel resolution, ensuring optimal capture of ultrastructural features during tilt-series acquisition. Dual-axis tilt series (ranging from - 60° to +60° or -68° to +68°) were acquired at 120 kV using SerialEM^83,84^ using Eagle CCD or Veleta CCD camera. 90° rotations between the two axes were manually performed. 6 out of 42 dual-axis tilt series acquisitions were made at Thermo Fisher Scientific’s Hillsboro NanoPort on a Talos L120C TEM at 120 kV using Thermo Scientific Tomo software.

### Tilt-series preprocessing and tomogram reconstruction

Tilt-series preprocessing and tomographic reconstruction were performed using IMOD^85^. All tilt series were denoised using IMOD’s mtffilter function with a low-pass cutoff of 0.166 and a Gaussian roll-off of 0.08 to suppress high-frequency noise. The tilt series were then downsampled to a 2 nm pixel resolution using IMOD’s squeezevol function to improve contrast. Using Etomo in IMOD, both axes were independently coarse-aligned using cross-correlation and fine-aligned using patch tracking or 10 nm gold fiducials, whenever fiducials were used. Tomograms for each axis were reconstructed using the simultaneous iterative reconstruction technique (SIRT) algorithm within Etomo for 20 iterations, with a cutoff of 0.4 and a Gaussian falloff of 0.08 for high-frequency noise suppression. Dual-axis tomograms were then combined using automatic patch fitting in Etomo’s tomogram combination panel. Combined dual-axis tomograms were flattened within Etomo’s post-processing panel to correct for warping caused by prolonged electron beam exposure during dual-axis tilt-series acquisitions.

The resulting tomograms measured approximately 1024 × 1024 × 50 voxels (XYZ) at an isotropic voxel size of 2 nm, yielding 2×2 × 0.1 μm³ volumes that captured partial 3D cross-sections of mitochondria. We tested the feasibility of increasing volumetric coverage and acquired two serial tomograms for one mitochondrion and a two-tile montage tomogram for another. Although it was technically feasible to acquire larger volumes of individual juxtaspinal mitochondrial regions, this approach introduced additional challenges during post-processing. We therefore limited subsequent acquisitions to single-section, single-tile volumes, prioritizing the sampling of a larger population of juxtaspinal mitochondrial regions to better capture biological variability rather than extending volume coverage of the same mitochondrion.

### Deep learning-based simultaneous segmentation of mitochondrial and surrounding structures

To segment mitochondrial compartments and nearby cellular structures simultaneously, we developed a custom volumetric segmentation network incorporating components of the 3D U-Net architecture, residual learning and bottleneck self-attention^86-88^. The model was trained to identify five structures in electron tomograms: the volume enclosed by the outer mitochondrial membrane (OMM), representing total mitochondrial volume; the volume enclosed by the inner boundary membrane (IBM), including the mitochondrial matrix and cristae lumen; the intermembrane space (IMS), including the peripheral intermembrane space and cristae lumen; the endoplasmic reticulum (ER); and ribosomes. These five labels were generated jointly by a single model in one inference pass rather than by separate feature-specific networks.

The labels were represented as separate, potentially overlapping binary channels to preserve their biological hierarchy. OMM was treated as the parent mitochondrial compartment, with IBM and IMS constrained to remain within OMM. ER and ribosomes were restricted to regions outside OMM. When ribosome and ER annotations overlapped because of their close spatial proximity, overlapping voxels were assigned to the ribosome class to preserve the ribosome annotation. Ribosomes were annotated only when they could be identified confidently within ribosome clusters. Ribosomes were classified according to their proximity to the OMM; however, OMM-proximal ribosomes were not further subdivided according to whether they were proximal to the OMM alone or to both the OMM and ER.

Model training was performed in two rounds. For the first round, all five structures were manually segmented in four tomograms using ORS Dragonfly (version 2022.2; Object Research Systems) to generate the initial ground-truth dataset. The first-round model was applied to eight additional tomograms, and its predictions were manually proofread in ORS Dragonfly to correct false-positive, false-negative and boundary errors. These corrected segmentations were added to the training dataset, producing a second-round training set of 12 tomograms. The second-round model was subsequently applied to the remaining tomograms. This iterative procedure increased the representation of mitochondrial morphologies, membrane orientations and image-quality conditions while reducing the amount of de novo manual annotation required.

Each tomogram was stored as an EM image stack with binary masks for OMM, IBM, IMS, ER and ribosomes. Ribosome annotations were unavailable for tomograms where no ribosomes were present. When a ribosome mask was unavailable, that channel was marked as missing and excluded from the associated loss and performance calculations rather than being interpreted as an empty segmentation. All available masks were required to have the same dimensions as the corresponding EM stack.

Each complete EM tomogram was independently z-score normalized by subtracting its mean intensity and dividing by its standard deviation. Normalized intensities were clipped to the range [−5, 5] to limit the influence of extreme voxel values while preserving relative contrast within each tomogram.

Tomograms were assigned to training, model-selection validation and independent test sets before patch extraction. The training, validation and test set contained 4, 1, 1 control and 4, 1, 1 homeostatic scaling tomograms from three independent preparations. No patch from a validation or test tomogram was used for training. Test tomograms were not used for model refinement, hyperparameter selection, checkpoint selection, probability-threshold selection, or qualitative monitoring. Reference segmentations for the test tomograms were generated manually and were not produced by correcting the evaluated model’s predictions.

The network was trained using patches measuring 32 × 256 × 256 voxels in Z, Y and X. Patches extending beyond the tomogram boundaries were padded with zeros. Because the five structures differed substantially in abundance, patch centers were sampled from voxels belonging to OMM, IBM, IMS, ER, or ribosome annotations and from unlabeled background and randomly selected locations. ER- and ribosome-centered patches were sampled more frequently because these structures were sparse. Class and positive-voxel weights were calculated from label prevalence in the training tomograms, with predefined limits preventing any single class from dominating the loss.

For on-the-fly data augmentation, training patches and their corresponding masks were randomly reflected along the X, Y and Z axes and rotated in 90° increments in the XY plane. EM intensities were randomly scaled and shifted, and Gaussian noise of variable magnitude was added. In a subset of patches, one to four consecutive Z slices were replaced with blank values to reduce sensitivity to localized information loss through the tomographic depth. Spatial transformations were applied identically to the EM images and masks, whereas intensity transformations were applied only to the EM images.

The encoder contained sequential three-dimensional residual blocks that progressively increased feature depth while reducing spatial resolution. Initial downsampling was restricted to the XY plane, followed by downsampling in all three dimensions at deeper encoder levels. This anisotropic design preserved information along the comparatively shallow Z dimension. The decoder restored the original spatial resolution using transposed convolutions and concatenated skip connections from the corresponding encoder levels.

A multi-head self-attention module was applied at the spatially compressed bottleneck. This allowed information from spatially separated regions of a patch to interact while limiting the computational cost of global attention. The decoded feature representation was passed to separate segmentation, boundary and affinity heads.

The segmentation head produced five probability maps using sigmoid activation. Hierarchical constraints were applied within the forward pass: IBM and IMS probabilities were multiplied by the predicted OMM probability; ER and ribosome probabilities were multiplied by the probability of being outside OMM; and ER probability was suppressed in regions with high ribosome probability. A hierarchy loss additionally penalized IBM or IMS predictions outside OMM, ER or ribosome predictions inside OMM, and overlap between ER and ribosome predictions.

The boundary head predicted the boundary of each of the five structures. Boundary targets were generated by identifying labeled-to-unlabeled transitions along the X, Y and Z axes. The affinity head predicted whether adjacent voxels along each axis belonged to the same OMM, IBM, IMS or ER structure. Ribosomes were excluded from the affinity task because they occurred as small punctate objects rather than continuous volumes.

The total loss combined weighted binary cross-entropy, Dice overlap, boundary classification, voxel affinity and hierarchy terms. The respective loss weights were 1.00, 1.00, 0.15, 0.15, and 0.20. Missing annotation channels were masked from all applicable loss terms. Training used 256 newly sampled patches per epoch, a batch size of one, and the AdamW optimizer with an initial learning rate of 2 × 10⁻⁴ and weight decay of 1 × 10⁻⁴. Training continued for up to 100 epochs. Final checkpoint selection was based on the mean class-specific Dice coefficient measured across the two complete validation tomograms. Final segmentation performance was measured on two independently held-out complete tomograms. Dice coefficient, precision and recall were calculated separately for OMM, IBM, IMS, ER and ribosomes. The resulting values were as follows for OMM, Dice 0.923, precision 0.883 and recall 0.967; IBM, Dice 0.912, precision 0.859 and recall 0.972; IMS, Dice 0.833, precision 0.734 and recall 0.965; ER, Dice 0.452, precision 0.329 and recall 0.730; and ribosomes, Dice 0.259, precision 0.348 and recall 0.206.

Training required ∼24 hours and was performed using PyTorch version 2.12.0 on an Apple M5 Max with 40 GPU cores and 128 GB unified memory using the Metal Performance Shaders backend. The final model generated all five segmentation channels simultaneously in ∼100 s per complete tomogram. Inference of the 42-tomogram dataset therefore required ∼70 min. All final predictions were inspected and proofread in ORS Dragonfly before three-dimensional reconstruction and morphometric analysis.

Published timing values were used only as contextual comparisons because hardware, tomogram dimensions, target structures, training data, and definitions of segmentation time differed between studies. Chen et al. trained and applied a separate binary network for each feature, whereas the present model generated five labels in one training run and one inference pass^54^. Heebner et al. subsequently demonstrated simultaneous segmentation of multiple cryo-ET structures and reported a total segmentation workflow of less than 30 min per tomogram in most cases^55^. Published studies have not reported proofreading person-hours separately, precluding a direct comparison of complete human–computer workflow time.

### 3D mesh processing and morphometric analysis of cristae, endoplasmic reticulum, and ribosomes

Mitochondria were captured at different orientations within the tomograms and could appear horizontally, vertically, or diagonally oriented in the field of view. The reconstructed volumes also varied in thickness because of differences in sectioning, sample shrinkage during prolonged electron beam exposure, and residual warping of the sections. Consequently, the amount of mitochondrial structure captured within the original tomogram boundaries was not directly comparable across samples. To ensure that equivalent mitochondrial regions were included in all downstream measurements, each tomogram and its corresponding segmentations were trimmed to encompass a standardized 2 μm juxtaspinal mitochondrial region. This normalization prevented differences in tomogram dimensions or mitochondrial orientation from producing unequal spatial sampling across mitochondrial regions.

Meshes for the IMS, IBM, OMM, endoplasmic reticulum (ER), ribosomes, and a bounding box representing the full tomogram volume were generated using the marching cubes algorithm in the scikit-image Python package^73,89^ applied to their respective segmentations via a custom Python script and saved as STL files. All meshes were then imported into the open-source platform Blender using the GAMer2 plugin. Within Blender, we applied the built-in ‘remesh’ and ‘triangulate’ functions to all meshes. Voxel-based remeshing was performed at a resolution of 2 nm for ribosomes and 4 nm for the IMS, IBM, OMM, and ER. Subsequent triangulation yielded lighter (i.e., reduced vertex count) and smoother meshes without compromising biological fidelity. All processed meshes were exported from Blender and saved as STL files.

At the initial stage, the IMS mesh included both the outer (OMM) and inner (IMM) mitochondrial membrane surfaces. In a fully reconstructed mitochondrion, these surfaces would naturally separate; however, since our reconstructions are partial, we implemented an alternative strategy to isolate the IMM. Using a custom Python script, we performed a Boolean subtraction of the IMS mesh from the IBM mesh. This operation produced the IMM mesh by incorporating cristae invaginations into the IBM surface while excluding the OMM. The resulting IMM mesh was imported into Blender, where the GAMer2 plugin was used to calculate the first-principal-curvature (k₁). The curvature values were exported from Blender as .pkl files, each containing a k₁ value for every vertex in the IMM mesh.

The k₁ values were imported into a custom Python script and assigned to the IMM mesh structure within the Python environment, where they were preserved throughout all downstream mesh operations. The IMM mesh included the IBM surface (from the original IBM mesh before Boolean subtraction), the CM surfaces, and the top and bottom capping surfaces corresponding to the start and end of the tomogram. These capping surfaces were required to maintain a closed, watertight mesh for volumetric Boolean operations. However, for surface area measurements, the capping surfaces needed to be removed. To achieve this removal, we used the bounding box mesh of the tomogram to identify and delete any IMM mesh vertices located within 5 nm of the bounding box surface, resulting in an open IMM surface mesh. To isolate the CM surface from the IBM, we used a custom Python function that referenced the original IBM mesh (before Boolean subtraction). Any region of the IMM mesh located more than 5 nm away from IBM mesh vertices was classified as CM, resulting in a CM surface mesh.

Crista junctions were determined automatically using the edges at the interface of CM and the rest of the IMM through a custom script. If these edges formed a ring, the centroid of the ring was marked as a point. Crista junction density was calculated as the number of points per total mitochondrial volume within the OMM mesh.

Using a Python script, we measured the CM surface area from the CM mesh, the IMM surface area from the IMM mesh, and the OMM volume (i.e., the total mitochondrial volume within the tomogram) from the OMM mesh. To quantify the surface area of highly curved cristae, we further extracted regions of the CM mesh with k₁ values greater than 80 μm^-1^. Using a Python script, we quantified ERMCS by separating the ER mesh within 20 nm of the OMM^90^ and measuring the surface area of the separated mesh, normalized to the total surface area of the ER. For quantifying ribosome density, we utilized the volume of the smallest single 3D mesh component of ribosomal clusters as unit ribosome volume and divided all cluster volumes by the unit volume. The ribosome cluster to OMM distance was measured as the shortest distance between the closest vertices of the ribosomal mesh to the OMM mesh.

### DNA-PAINT imaging and analysis

Rat primary hippocampal neurons were transfected with OMM-EGFP and PSD95.FingR-mTagBFP2 (PSD95FingR-BFP) between DIV 15-17 to label mitochondria and spines, respectively. At DIV18–19, neurons were treated with 2 μM tetrodotoxin (TTX) or maintained under control conditions. Neurons were fixed 24 hours after treatment. Fixation was performed using 4% paraformaldehyde (PFA) prepared in E4 imaging medium for 10 min at room temperature. Samples were then washed three times for 5 min in phosphate-buffered saline (PBS, pH 7.4).

Fixed neurons were permeabilized with 0.25% Triton X-100 in PBS for 20 min at room temperature and washed twice for 5 min in PBS. Samples were blocked with 4% bovine serum albumin (BSA) in PBS for 1 hour at room temperature. ATP synthase was labeled using a mouse monoclonal antibody against the OSCP subunit (OSCP Antibody A-8; Santa Cruz Biotechnology, sc-365162), and MIC60 was labeled using a rabbit polyclonal antibody against IMMT (Proteintech, 10179-1-AP). Primary antibodies were diluted in antibody incubation buffer (Massive Photonics) to final concentrations of 5 μg/ml for OSCP and 1.2 μg/ml for MIC60. Neurons were incubated with both primary antibodies overnight at 4°C. Samples were then washed three times for 5 min in PBS. Anti-mouse and anti-rabbit single-domain secondary antibodies carrying a distinct single-stranded DNA docking strand for sequential, multiplexed imaging (secondary dockers, sdAB) (Massive Photonics) were each diluted 1:400 in antibody incubation buffer (Massive Photonics) and applied simultaneously for 1 hour at room temperature. The anti-mouse sdAB docker was used to label OSCP, whereas the anti-rabbit sdAB docker was used to label MIC60. Samples were subsequently washed three times for 5 min in PBS.

To stabilize the antibody complexes, neurons were post-fixed with 4% PFA for 5 min at room temperature and washed three times for 5 min in PBS. To facilitate drift correction during downstream analysis, samples were then incubated with 90 nm nanogold fiducial markers (NanoPartz) diluted 1:50 for 20 min, followed by two rapid washes in PBS. To reduce nonspecific DNA interactions, neurons were incubated with salmon sperm DNA diluted 1:200 for 30 min and washed twice rapidly in PBS. FAST DNA-PAINT imagers were then added at final dilutions of 1:4000 for the F1 imager and 1:1000 for the F2 imager (Massive Photonics). Dye-labeled imager strands transiently hybridize to their complementary docking strands, producing the repeated ‘blinking’ from which single molecules were localized.

Neurons expressing OMM-EGFP were identified, and multiple imaging locations were selected for overnight acquisition. Three-dimensional DNA-PAINT imaging was performed using biplane acquisition on a Vutara VXL super-resolution microscope (Bruker). MIC60 and ATP synthase were imaged simultaneously using the 561 and 647 nm channels, respectively. Each acquisition cycle consisted of 1,200 simultaneous 561 and 647 nm frames, followed by an intermittent 488 nm OMM-EGFP reference frame. This sequence was repeated for 50 cycles, yielding a total of 60,000 frames for molecular localization and 50 OMM-EGFP reference frames for monitoring the mitochondrial position throughout the acquisition.

At the end of each DNA-PAINT acquisition, a three-dimensional Z-stack of OMM-EGFP-labeled mitochondria and PSD95FingR-BFP-labeled postsynaptic densities was acquired. The 3D Z-stack OMM-EGFP channel provided a wide Z-range of mitochondrial reference, and the PSD95FingR-BFP channel identified the positions of dendritic spines. The intermittent OMM-EGFP reference images collected during biplane imaging allowed mitochondrial position to be related to the terminal three-dimensional reference stack.

Single-molecule localizations were calculated using Bruker SRX software with experimentally calibrated three-dimensional point spread functions. Localizations positioned more than 450 nm above or below the focal plane were removed, restricting the analysis to a 900 nm axial range in which three-dimensional localization could be performed reliably. Lateral drift correction in XY was performed using redundant cross-correlation in Picasso, whereas axial drift correction in Z was performed using a custom script.

The three-dimensional reference stacks containing the mitochondrial and spine channels were deconvolved and converted into maximum-intensity projections. Mitochondria in the deconvolved OMM-EGFP image were registered to the OMM-EGFP reference images acquired during biplane imaging. The same spatial transformation was then applied to the PSD95FingR-BFP channel. This procedure placed the spine reference and the three-dimensional molecular localizations within the same coordinate system, allowing ATP synthase and MIC60 copies to be assigned according to their position relative to individual mitochondria and nearby dendritic spines.

Juxtaspinal and non-juxtaspinal mitochondrial regions were manually identified from the registered, deconvolved mitochondrial and spine images using the Time Series Analyzer plugin in Fiji. The coordinates of the manually defined regions were exported for subsequent assignment and quantification of ATP synthase and MIC60 copies.

Repeated DNA-PAINT localizations originating from individual molecular targets were grouped using the SMLM Clusterer in Picasso. Clustering was performed separately for the ATP synthase and MIC60 channels using a minimum of 10 localizations per cluster. Because localization precision varied between fields of view and channels, the clustering distances were determined from the corresponding nearest-neighbor-based localization precision (NeNA). The XY clustering radius was set to twice the NeNA value, and the Z clustering radius was set to four times the NeNA value for each molecular channel within each field of view.

Each resulting localization cluster was considered to represent a single labeled copy, and the three-dimensional centroid of the cluster was used as its position. The coordinates of the manually marked juxtaspinal and non-juxtaspinal mitochondrial regions were then used to identify cluster centroids located within each region. The numbers and spatial distributions of ATP synthase and MIC60 copies were then quantified within juxtaspinal and non-juxtaspinal mitochondrial regions.

### Phos-tag Mobility Shift Assay

Neurons were transfected with OMM-EGFP and Matrix-PKI or IMS-PKI on DIV 15-16. On DIV 18-19, neurons were treated with 2 μM TTX for 24 hours. Neurons were then lysed in mitochondrial isolation buffer (210 mM mannitol, 70 mM sucrose, 5 mM HEPES, 1 mM EGTA, 0.5% BSA, supplemented with protease inhibitors (Pierce, A32953) and phosphatase inhibitors (Pierce, A32957)), homogenized using a Dounce homogenizer, and centrifuged at 800 x g for 10 min at 4°C. To selectively measure phosphorylated levels of MIC60 in transfected neurons, mitochondria were immunoprecipitated from the resulting supernatant using GFP-Trap agarose beads (ChromoTek, gta). Supernatant was incubated with GFP-Trap beads for 2 hours at 4 °C, followed by three washes (5 min each) in mitochondrial isolation buffer to remove unbound material. Mitochondrial proteins were eluted by adding an equal volume of 2x Laemmli sample buffer to the beads and heating at 95 °C for 10 min.

For the phos-tag mobility shift assay, samples were run on 12.5% SuperSep Phos-tag gels (50 μmol/L) at 30 mA for 2-3 hours. Gels were then equilibrated (twice for 10 minutes each) in transfer buffer containing 10 mM EDTA, followed by a wash in transfer buffer without EDTA, before transfer onto a PVDF membrane at 400 mA for 2 hours. Membranes were immunoblotted for anti-MIC60 (1:1000) (Proteintech, 10179-1-AP) to resolve phosphorylated and non-phosphorylated species by mobility shift, and anti-GFP (1:1000) (Abcam, Ab290) to confirm immunoprecipitation of OMM-EGFP.

### Phospho/Western Blots

Total neuronal lysate samples, reserved from the same lysate prepared for the phos-tag mobility shift assay above, were run for standard SDS-PAGE on mPAGE 10% Bis-Tris Precast gels (Millipore) at 120-150 V for 1-2 hours and transferred onto PVDF membrane at 400 mA for 90 minutes. Membranes were probed using the following antibodies: anti-MIC60 (1:1000) (Proteintech, 10179-1-AP), anti-GFP (1:1000) (Abcam, Ab290), anti-GluA1 (1:1000) (Millipore Sigma, SAB5201086), anti-phospho-GluA1 S845 (1:1000) (Cell Signaling, 8084T), anti-GluN1 (1:1000) (Cell Signaling, 5704T), and anti-phospho-GluN1 S897 (1:1000) (Cell Signaling, 3385S). Blots were imaged using Amersham ImageQuant 800, and band intensities were quantified using Image Studio Lite. The raw intensity value of a phosphorylated protein was divided by the raw intensity value of the corresponding total protein to obtain a phosphorylated-to-total protein ratio.

### Quantification and statistical analysis

All statistical analyses were performed using R (v4.3.0). Measurements made per spine or mitochondrion were averaged per neuron when the number of measurements exceeded several hundred. The effect of experimental condition was assessed using two-way ANOVA with condition and animal as factors. When both experimental condition and mito region were compared, three-way ANOVA was performed with condition, mito region, and biological replicates as factors. When a significant effect involving condition was detected, Tukey-adjusted post hoc comparisons were used to compare experimental groups. The Kruskal–Wallis test was used for nonparametric comparisons. One-way repeated-measures ANOVA was used for time-course data. Western blot data were analyzed using a matched two-way ANOVA, with construct and TTX treatment as factors and biological replicates (gels) as the matched block, since all conditions for each biological replicate were analyzed in parallel on the same gel and are not statistically independent^91,92^, following the two-way repeated-measures framework of Maxwell and Delaney (2004, Ch. 12). This approach is consistent with prior applications of repeated-measures ANOVA to Western blot densitometry, including analyses of phospho/total protein ratios across parallel biological samples and comparisons of treatment groups analyzed on the same blot^93,94^. Electrophysiological event distributions were compared using two-sample Kolmogorov–Smirnov tests. Relationships between continuous variables in the connectomics analyses were evaluated using Pearson correlation, with linear regression lines used to display the fitted relationships.

Statistical details of experiments, including statistical tests and n and p values, are mentioned in the figure legends. Each biological replicate corresponds to one weekly batch of neuronal culture preparation. Data visualizations were generated using ggplot2 (v3.5.0) and ggbeeswarm (v0.7.2). In all the figures, the box whisker plots represent the median (line), mean (square point), 25th-75th percentile, i.e. interquartile range (box), 1.5xinterquartile range (whisker), and individual data points (solid circles shaded by biological replicates). For regression models, fixed-effect prediction lines and confidence ribbons were computed manually from model coefficients and variance-covariance matrices. A p-value of less than or equal to 0.05 was considered significant for all statistical tests. No statistical method was used to predetermine the sample size. Sample sizes were similar to or larger than those reported in the previous publications in the field and sufficient for our claims based on statistical significance.

### Data and materials availability

Plasmids generated in this study will be deposited in Addgene. Source data are provided with this paper. Custom-written scripts for data analysis, quantification, and statistics, along with demo images and user instructions, are available at https://github.com/Rangaraju-Lab/Shah-et-al-2026. Requests for further information and resources should be directed to and will be fulfilled by Vidhya Rangaraju.

