## Supplemental figures and legends for "Mitochondria remodel near synapses to boost metabolic capacity and fuel plasticity"

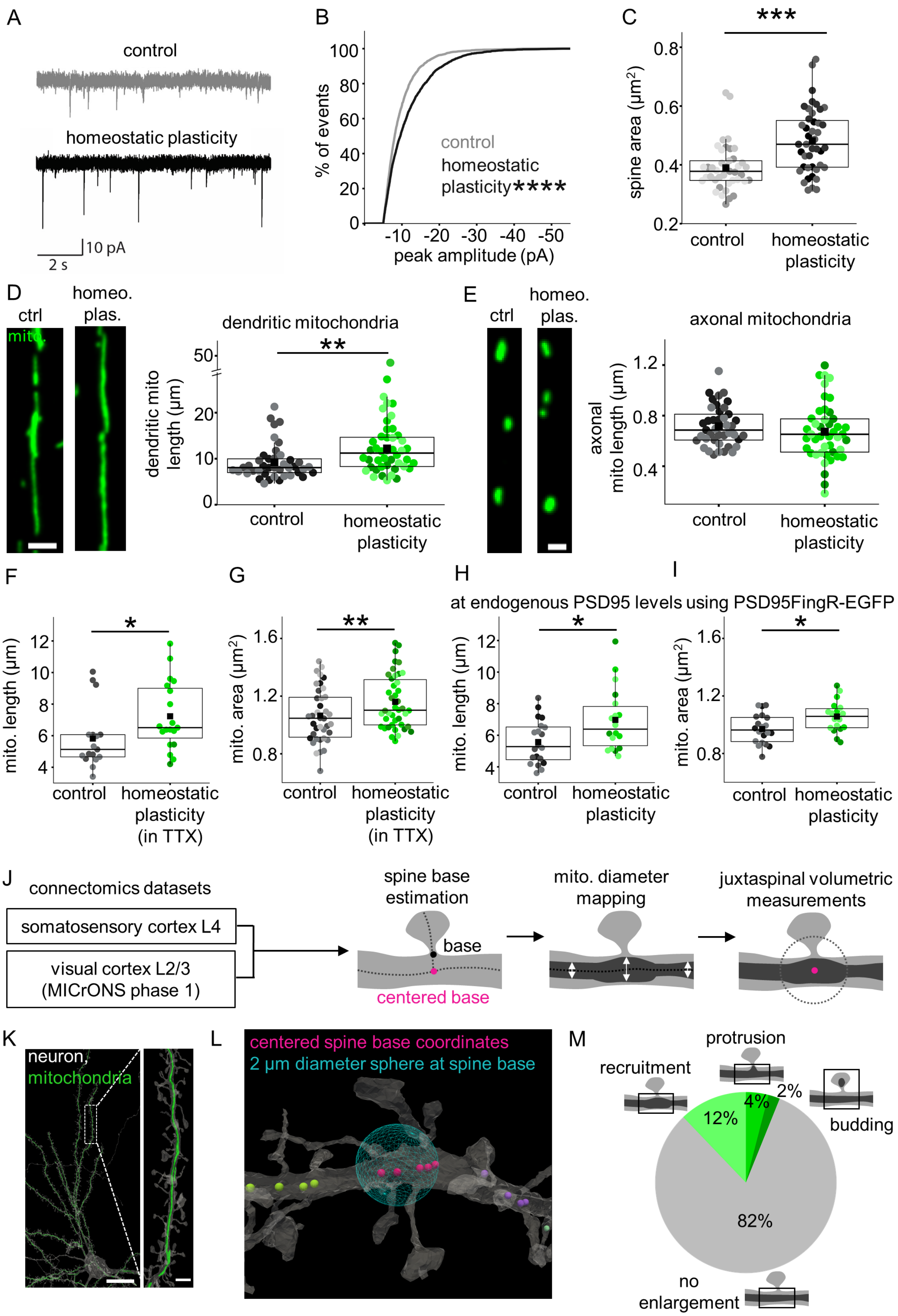
A
control
homeostatic plasticity
10 pA
2 s
B
% of events
control
homeostatic plasticity****
peak amplitude (pA)
C
***
spine area (µm²)
control
homeostatic plasticity
D
ctrl
homeo. plas.
mito.
dendritic mitochondria
**
dendritic mito length (µm)
control
homeostatic plasticity
E
ctrl
homeo. plas.
axonal mitochondria
axonal mito length (µm)
control
homeostatic plasticity
F
*
mito. length (µm)
control
homeostatic plasticity (in TTX)
G
**
mito. area (µm²)
control
homeostatic plasticity (in TTX)
H
at endogenous PSD95 levels using PSD95FingR-EGFP
*
mito. length (µm)
control
homeostatic plasticity
I
*
mito. area (µm²)
control
homeostatic plasticity
J
connectomics datasets
somatosensory cortex L4
visual cortex L2/3 (MICrONS phase 1)
spine base estimation
base
centered base
mito. diameter mapping
juxtaspinal volumetric measurements
K
neuron, mitochondria
L
centered spine base coordinates
2 µm diameter sphere at spine base
M
protrusion
recruitment
budding
12%
4%
2%
82%
no enlargement

**Figure S1. Validation and control experiments related to juxtaspinal mitochondrial enlargement.**

**A** Representative electrophysiological recordings (quantified in **B**) of miniature excitatory postsynaptic currents (mEPSCs) in control and homeostatic plasticity-induced neurons. **B** Cumulative distribution of mEPSC peak amplitude shows a significant increase in homeostatic plasticity-induced neurons (black) relative to control (gray). n in neurons, biol. replicates: 10, 5 (control), 12, 5 (homeostatic plasticity). Kolmogorov–Smirnov test, p-value: <0.0001. **C** Average spine area increases in homeostatic plasticity-induced neurons (black) relative to control (gray). Different shades within each condition denote independent biological replicates. n in neurons, biol. replicates: 41, 3 (control), 45, 3 (homeostatic plasticity). Two-way ANOVA (condition*biol. replicates), p-value: <0.0001. **D**, **E** Average mitochondrial length increases in homeostatic plasticity-induced neurons (green) relative to control (gray) in dendrites (**D**), but not in axons (**E**). n in neurons, biol. replicates: 41, 3 (dendrites, control), 45, 3 (dendrites, homeostatic plasticity), 41, 3 (axons, control), 45, 3 (axons, homeostatic plasticity). Two-way ANOVA (condition*biol. replicates), p-values: 0.0029 (dendrites: control vs. homeostatic plasticity), 0.3186 (axons: control vs. homeostatic plasticity). Scale bars: 5 µm (dendrites), 1 µm (axons). **F** Average dendritic mitochondrial length increases in homeostatic plasticity-induced neurons imaged in the continued presence of TTX (green) relative to control (gray). n in neurons, biol. replicates: 17, 2 (control), 19, 2 (homeostatic plasticity in TTX). Two-way ANOVA (condition*biol. replicates), p-value: 0.0357. **G** Average juxtaspinal mitochondrial area increases in homeostatic plasticity-induced neurons imaged in the continued presence of TTX (green) relative to control (gray). n in neurons, biol. replicates: 38, 5 (control), 42, 5 (homeostatic plasticity in TTX). Two-way ANOVA (condition*biol. replicates), p-value: 0.009. **H** Average dendritic mitochondrial length increases in homeostatic plasticity-induced neurons (green) relative to control (gray) at endogenous PSD95 levels, detected using PSD95FingR-EGFP. n in neurons, biol. replicates: 20, 3 (control), 18, 3 (homeostatic plasticity). Two-way ANOVA (condition*biol. replicates), p-value: 0.0202. **I** Average juxtaspinal mitochondrial area increases in homeostatic plasticity-induced neurons (green) relative to control (gray) at endogenous PSD95 levels, detected using PSD95FingR-EGFP. n in neurons, biol. replicates: 20, 3 (control), 18, 3 (homeostatic plasticity). Two-way ANOVA (condition*biol. replicates), p-value: 0.0176. **J** Data-analysis workflow for quantifying juxtaspinal mitochondrial widening in the mouse somatosensory and visual cortex connectomics datasets. **K** Representative 3D rendering (related to **Fig. 1H-J**) of a neuron (gray) with dendritic mitochondria (green) from the visual cortex L2/3 MICrONS phase 1 connectomics dataset. Scale bar: 20 µm. Inset, dendritic segment with spines (gray) and mitochondria (green). Scale bar: 2 µm. **L** Representative 3D rendering of a dendritic segment (related to **Fig. 1I, J**) showing spine-base centers (green, pink, and purple dots denote different spine clusters; see Methods) and a 2 µm sphere around the center of one spine cluster (blue) used to quantify juxtaspinal mitochondrial volume. **M** Fraction of juxtaspinal mitochondrial regions exhibiting recruitment, protrusion, budding, and no enlargement (analogous to no change from **Fig. 1C**) observed as static occurrences in somatosensory cortex, n in mito. reg., 24081 (recruitment), 7990 (protrusion), 4113 (budding), 162379 (no enlargement/no change).

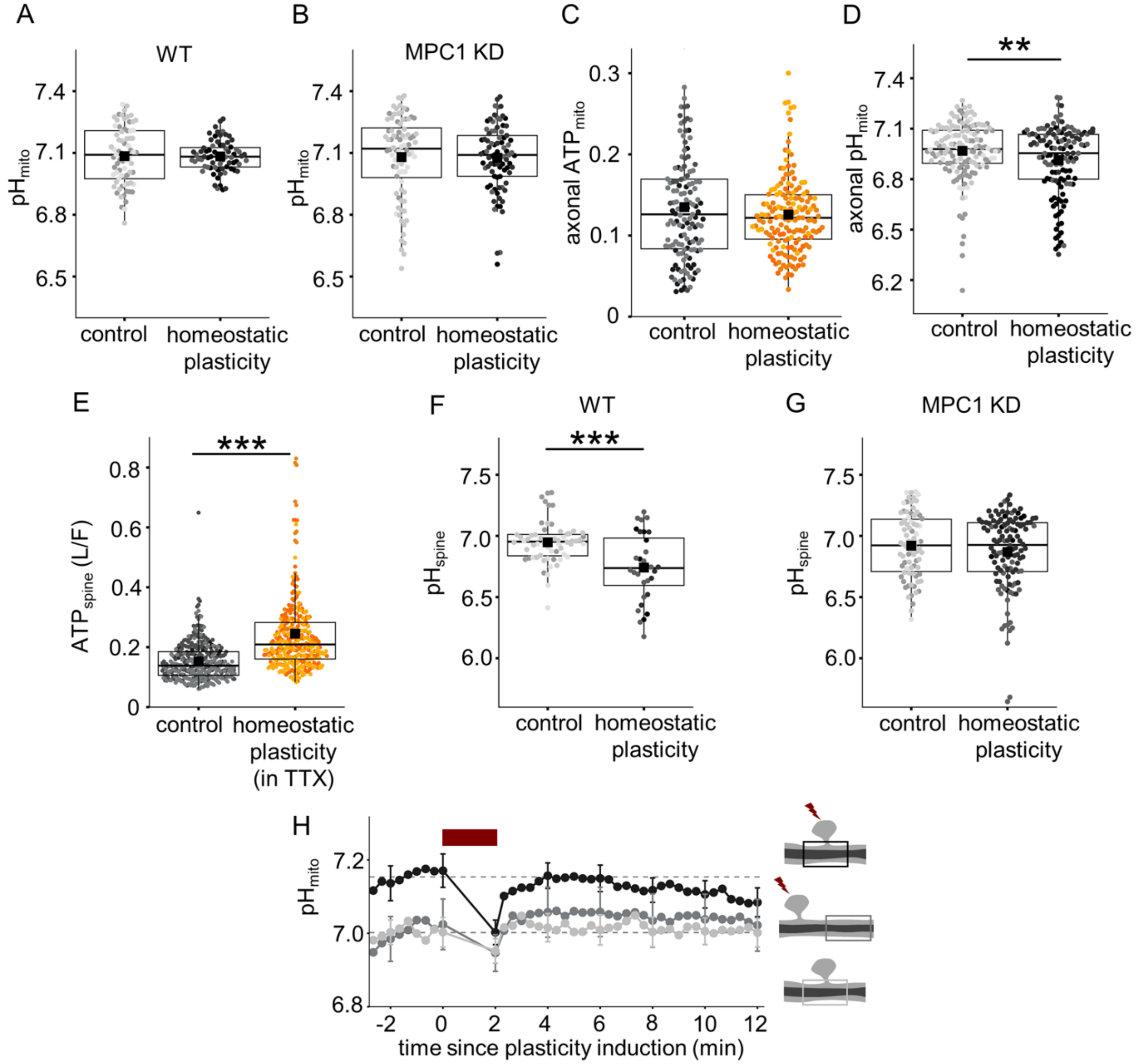

**Figure S2. Control experiments related to $ATP_{mito}$ and $ATP_{spine}$ measurements**

**A, B** Average $pH_{mito}$ is unchanged between control (gray) and homeostatic plasticity-induced neurons (dark gray) in WT (**A**) and MPC1 KD (**B**). Different shades within each condition denote independent biological replicates. n in mitochondria, biol. replicates: 73, 3 (WT control), 71, 3 (WT homeostatic plasticity), 84, 3 (MPC1 KD control), 85, 3 (MPC1 KD homeostatic plasticity). Two-way ANOVA (condition*biol. replicates), p-value: 0.9496 (WT), 0.8888 (MPC1 KD). **C** In axons, average $ATP_{mito}$ is unchanged between control (gray) and homeostatic plasticity-induced neurons (dark gray). n in mitochondria, biol. replicates: 135, 3 (control), 161, 3 (homeostatic plasticity). Two-way ANOVA (condition*biol. replicates), p-value: 0.1213. **D** In axons, average $pH_{mito}$ showed a small but significant decrease in homeostatic plasticity-induced neurons (dark gray) relative to control (gray). n in mitochondria, biol. replicates: 141, 3 (control), 129, 3 (homeostatic plasticity). Two-way ANOVA (condition*biol. replicates), p-value: 0.007. **E** Average $ATP_{spine}$ measured in the continued presence of TTX increases in homeostatic plasticity-induced neurons (orange) relative to control (gray). The control dataset is the same as in **Fig. 2H**. n in spines, biol. replicates: 343, 3 (control), 329, 3 (homeostatic plasticity). Two-way ANOVA (condition*biol. replicates), p-value: <0.0001. **F, G** Average $pH_{spine}$ decreases in homeostatic plasticity-induced neurons (dark gray) relative to control (gray) in WT (**F**), with no change in MPC1 KD (**G**). n in spines, biol. replicates: 60, 3 (WT control), 32, 3 (WT homeostatic plasticity), 78, 3 (MPC1 KD control), 120, 3 (MPC1 KD homeostatic plasticity). Two-way ANOVA (condition*biol. replicates), p-values: <0.0001 (WT), 0.1661 (MPC1 KD). **H** Average time course of $pH_{mito}$ in juxtaspinal mitochondrial regions (black), non-juxtaspinal mitochondrial regions (gray), and mitochondrial regions near uninduced spines (light gray) upon single-spine plasticity induction (red bar, 0.5 Hz, 120s) used to correct $ATP_{mito}$ in **Fig. 2K**. n in mito. reg., biol. replicates: 14, 7 (juxtaspinal mito. reg.), 11, 6 (non-juxtaspinal mito. reg.), 16, 4 (uninduced mito. reg.).

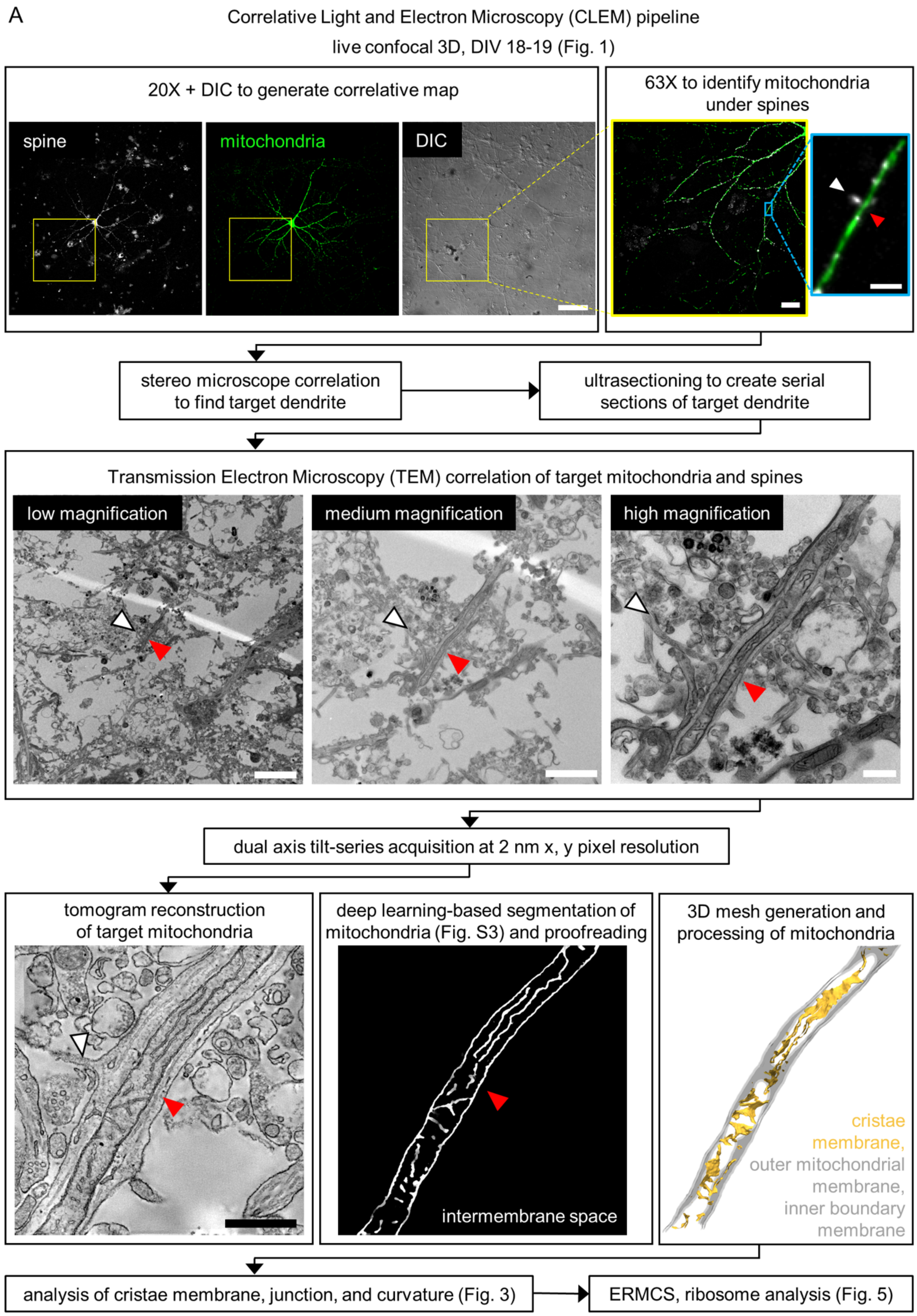

A
Correlative Light and Electron Microscopy (CLEM) pipeline
live confocal 3D, DIV 18-19 (Fig. 1)
20X + DIC to generate correlative map
spine
mitochondria
DIC
63X to identify mitochondria under spines
stereo microscope correlation to find target dendrite
ultrasectioning to create serial sections of target dendrite
Transmission Electron Microscopy (TEM) correlation of target mitochondria and spines
low magnification
medium magnification
high magnification
dual axis tilt-series acquisition at 2 nm x, y pixel resolution
tomogram reconstruction of target mitochondria
deep learning-based segmentation of mitochondria (Fig. S3) and proofreading
intermembrane space
3D mesh generation and processing of mitochondria
cristae membrane,
outer mitochondrial membrane,
inner boundary membrane
analysis of cristae membrane, junction, and curvature (Fig. 3)
ERMCS, ribosome analysis (Fig. 5)

**Figure S3. Correlative light and electron microscopy pipeline used to identify and analyze juxtaspinal mitochondria.**

Schematic of the correlative light and electron microscopy workflow in primary hippocampal neuronal cultures. Dendritic spines (white arrowheads) and juxtaspinal mitochondria (red arrowheads) were first targeted by fluorescence microscopy and differential interference contrast imaging (DIC) at 20x and 63x magnification, then relocated in transmission electron microscopy images (white and red arrowheads, TEM correlation) for tomogram acquisition, reconstruction, deep-learning-based segmentation, and proofreading. Scale bars: 100 µm (20x confocal), 20 µm (63x confocal), 2 µm (63x confocal inset), 5 µm (low-magnification TEM), 2 µm (medium-magnification TEM), 0.5 µm (high-magnification TEM), 0.5 µm (tomogram). The reconstructed juxtaspinal mitochondrial mesh and tomogram shown are Mitochondrion 4 of **Fig. 3B, C, S5**.

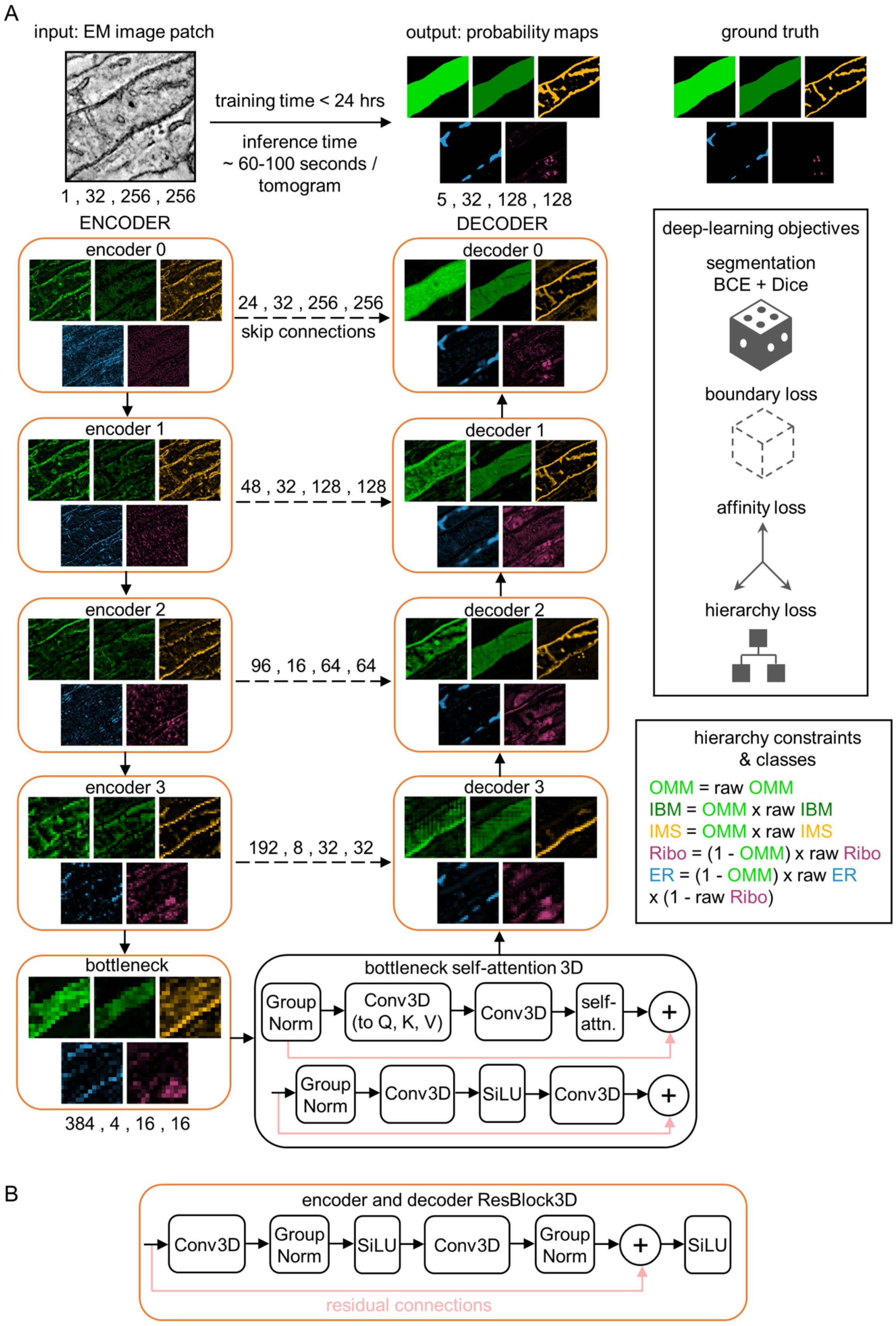
A
input: EM image patch
1 , 32 , 256 , 256
training time < 24 hrs
inference time
~ 60-100 seconds / tomogram
output: probability maps
5 , 32 , 128 , 128
ground truth
ENCODER
DECODER
encoder 0
encoder 1
encoder 2
encoder 3
bottleneck
384 , 4 , 16 , 16
decoder 0
decoder 1
decoder 2
decoder 3
24 , 32 , 256 , 256
skip connections
48 , 32 , 128 , 128
96 , 16 , 64 , 64
192 , 8 , 32 , 32
deep-learning objectives
segmentation
BCE + Dice
boundary loss
affinity loss
hierarchy loss
hierarchy constraints & classes
OMM = raw OMM
IBM = OMM x raw IBM
IMS = OMM x raw IMS
Ribo = (1 - OMM) x raw Ribo
ER = (1 - OMM) x raw ER x (1 - raw Ribo)
bottleneck self-attention 3D
Group Norm
Conv3D (to Q, K, V)
Conv3D
self-attn.
+
Group Norm
Conv3D
SiLU
Conv3D
+
B
encoder and decoder ResBlock3D
Conv3D
Group Norm
SiLU
Conv3D
Group Norm
+
SiLU
residual connections

88

**Figure S4. Hierarchy-constrained 3D residual U-Net with self-attention for simultaneous segmentation of mitochondrial and surrounding structures**

**A** Schematic of the deep-learning architecture used for simultaneous segmentation of the mitochondrial outer membrane-enclosed volume (OMM), inner boundary membrane-enclosed volume (IBM), intermembrane space (IMS), endoplasmic reticulum (ER), and ribosomes (related to **Fig. 2 and 4**). Input: a representative slice from a small 3D electron tomogram patch. The encoder consists of sequential residual blocks that progressively increase feature depth while reducing spatial resolution, initially within the XY plane and subsequently in all three dimensions. A self-attention module at the bottleneck integrates spatial information across the complete patch. The decoder restores spatial resolution using upsampling and concatenated skip connections that transfer features from the corresponding encoder levels. **B** Computation performed within each residual block, outlined in orange. Representative feature maps that best capture each of the five segmented structures are shown across the encoder and decoder to illustrate progressive refinement of structural information. Output: probability maps for the OMM (green), IBM (dark green), IMS (orange), ER (blue), and ribosomes (magenta); these colors denote model output channels and differ from the rendering colors used in **Fig. 2 and 4**. Learning objectives combine binary cross-entropy (BCE), Dice, affinity, and hierarchy losses to optimize segmentation accuracy, boundary relationships, and contextual organization across organelles. Biological hierarchy constraints restrict IBM and IMS predictions to the OMM volume and ER and ribosome predictions to regions outside mitochondria, while suppressing overlap between ER and ribosomes.

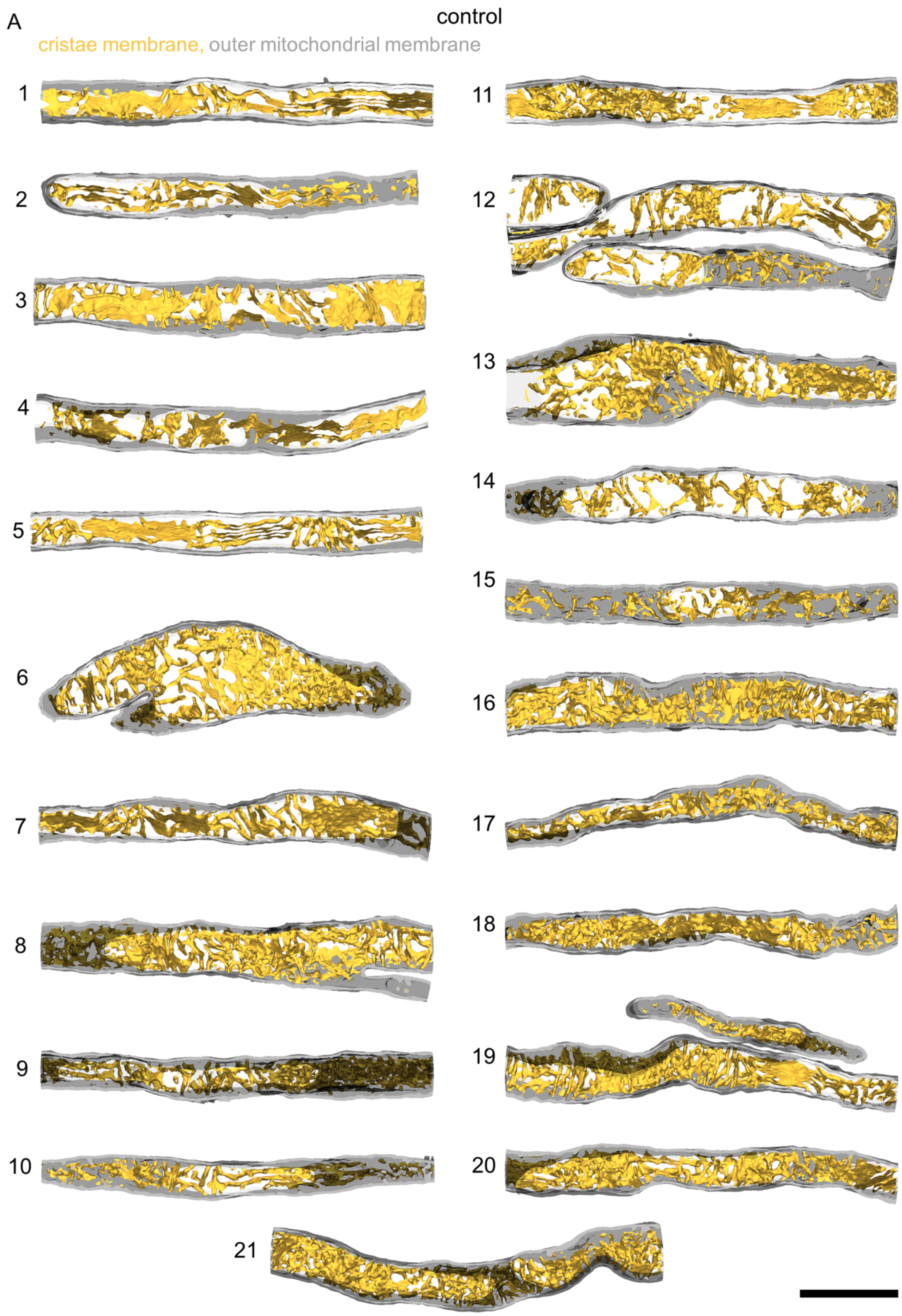
A
control
cristae membrane, outer mitochondrial membrane

111

**Figure S5. Reconstructed electron tomograms showing cristae membrane in control.**

Reconstructed electron tomograms of juxtaspinal mitochondrial regions in control neurons (related to **Fig. 3C-H**), showing cristae membrane (yellow) and outer mitochondrial membrane (gray). n in reconstructions, biol. replicates: 21, 3. Mitochondrion 4 is the same as in **Fig. 3C**. Scale bar: 0.5 µm.

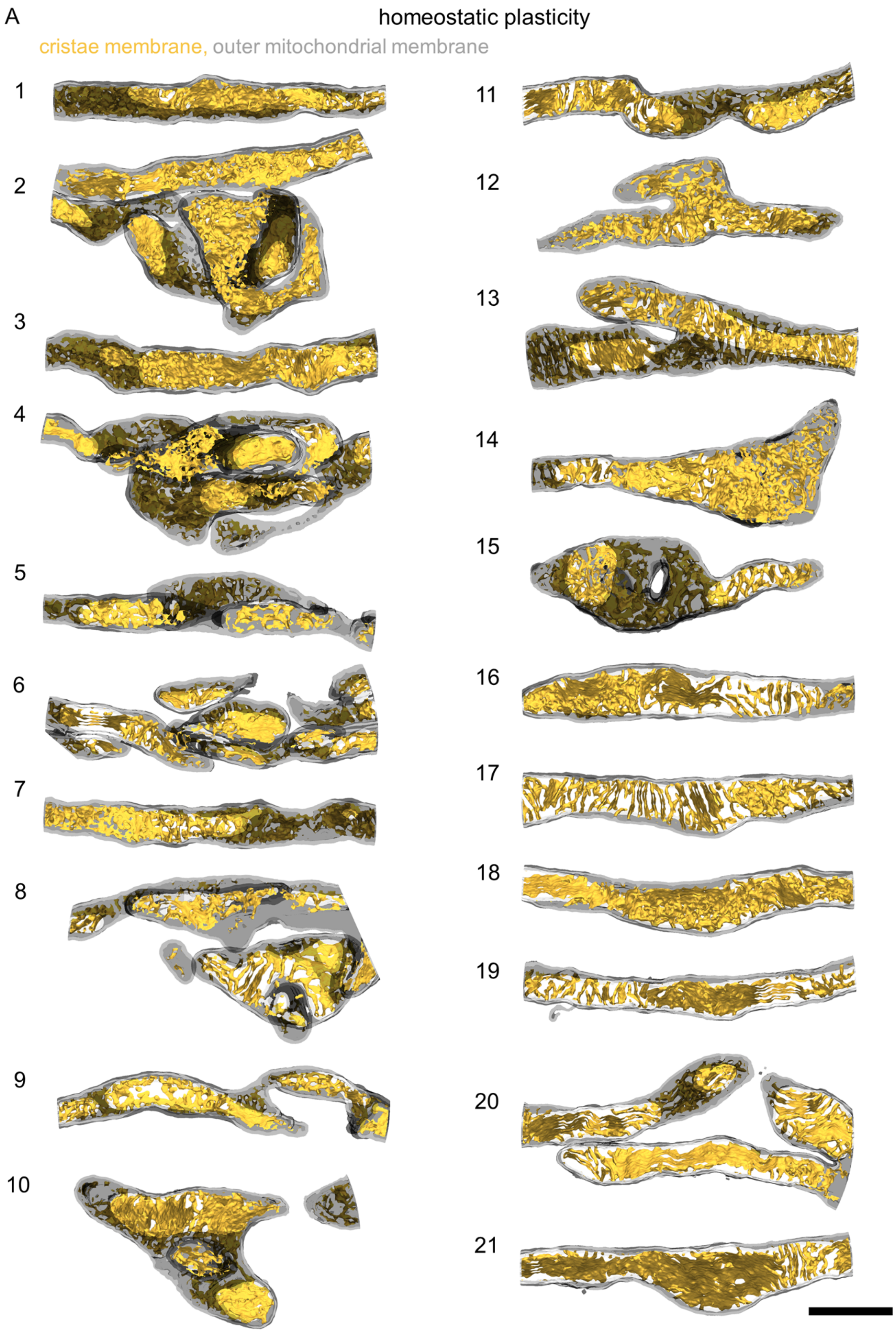
A
homeostatic plasticity
cristae membrane, outer mitochondrial membrane

118

**Figure S6. Reconstructed electron tomograms showing increased cristae membrane following synaptic plasticity.**

Reconstructed electron tomograms of juxtaspinal mitochondrial regions following homeostatic plasticity (related to **Fig. 3C-H**), showing cristae membrane (yellow) and outer mitochondrial membrane (gray)**.** n in reconstructions, biol. replicates: 21, 3. Mitochondrion 8 is the same as in **Fig. 3C**. Scale bar: 0.5 µm.

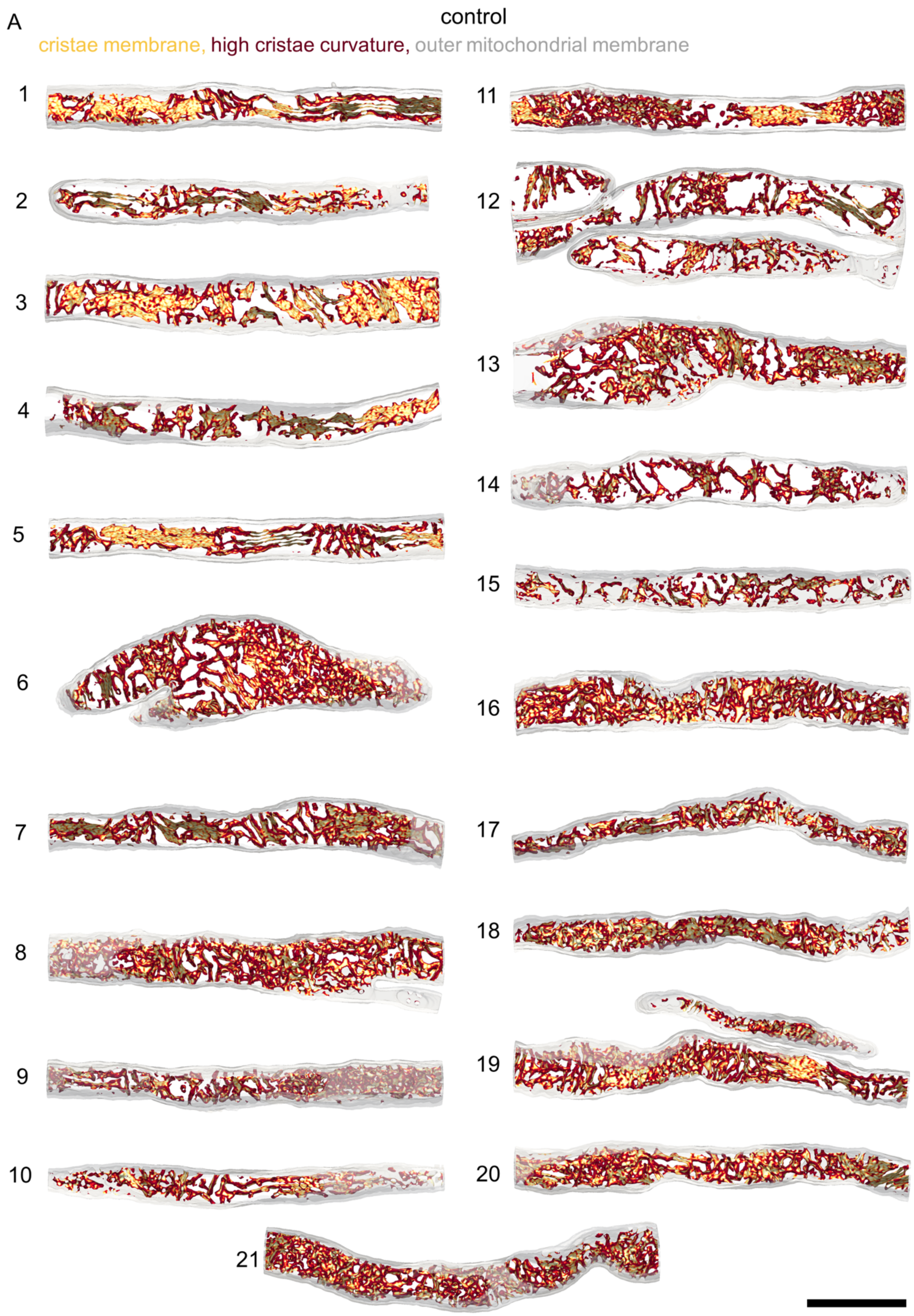
A
control
cristae membrane, high cristae curvature, outer mitochondrial membrane

125

**Figure S7. Reconstructed electron tomograms showing high cristae curvature in control.**

Reconstructed electron tomograms of juxtaspinal mitochondrial regions in control neurons (related to **Fig. 3G, H**), showing cristae membrane (yellow) and high first-principal-curvature ($k_1$) cristae membrane (red)**.** n in reconstructions, biol. replicates: 21, 3. Mitochondrion 4 is the same as in **Fig. 3G**. Scale bar: 0.5 µm.

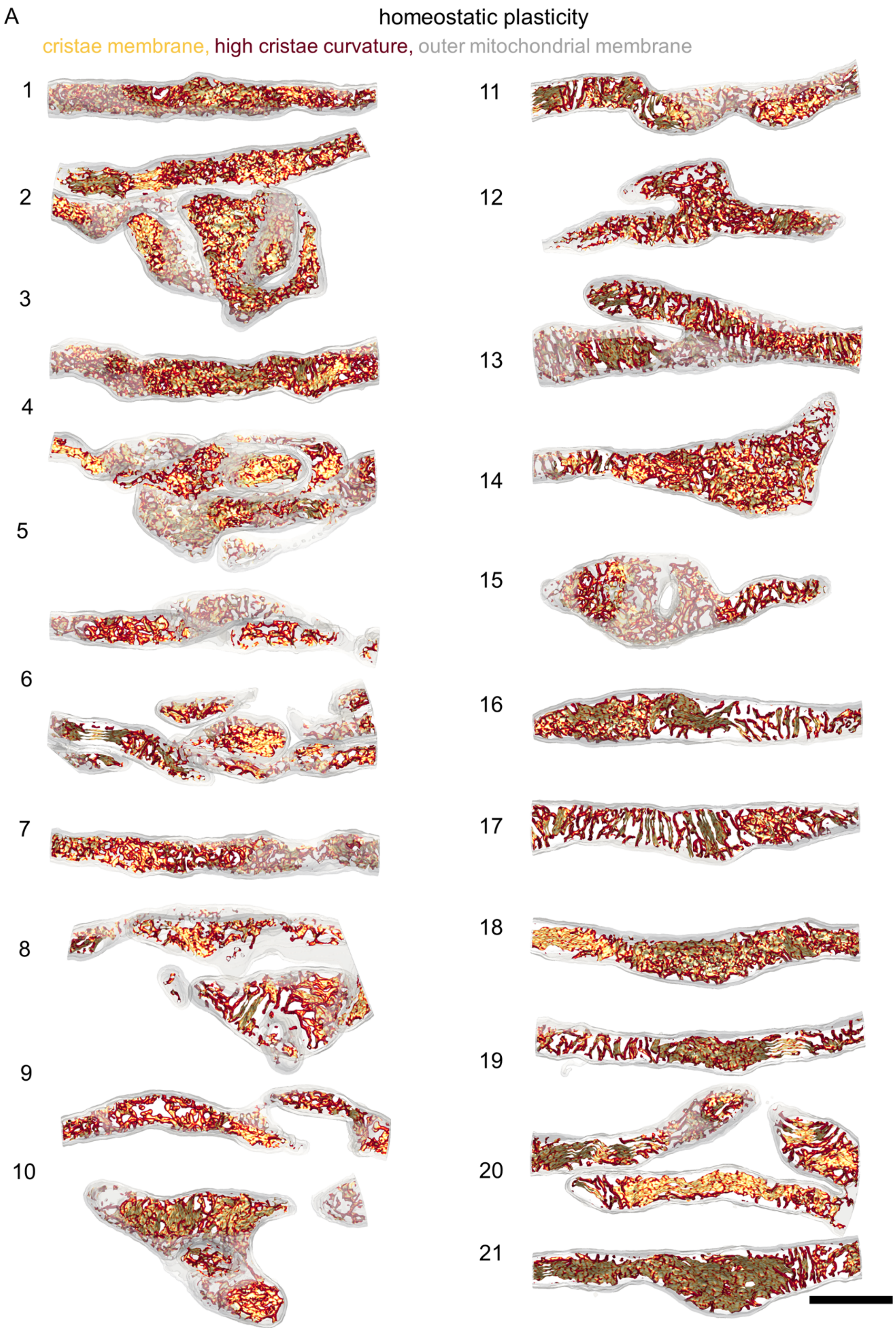
A
homeostatic plasticity
cristae membrane, high cristae curvature, outer mitochondrial membrane

132

**Figure S8. Reconstructed electron tomograms showing increased high cristae curvature following synaptic plasticity.**

Reconstructed electron tomograms of juxtaspinal mitochondrial regions following homeostatic plasticity (related to **Fig. 3G, H**), showing cristae membrane (yellow) and high first-principal-curvature ($k_1$) cristae membrane (red)**.** n in reconstructions, biol. replicates: 21, 3. Mitochondrion 8 is the same as in **Fig. 3G**. Scale bar 0.5 µm.

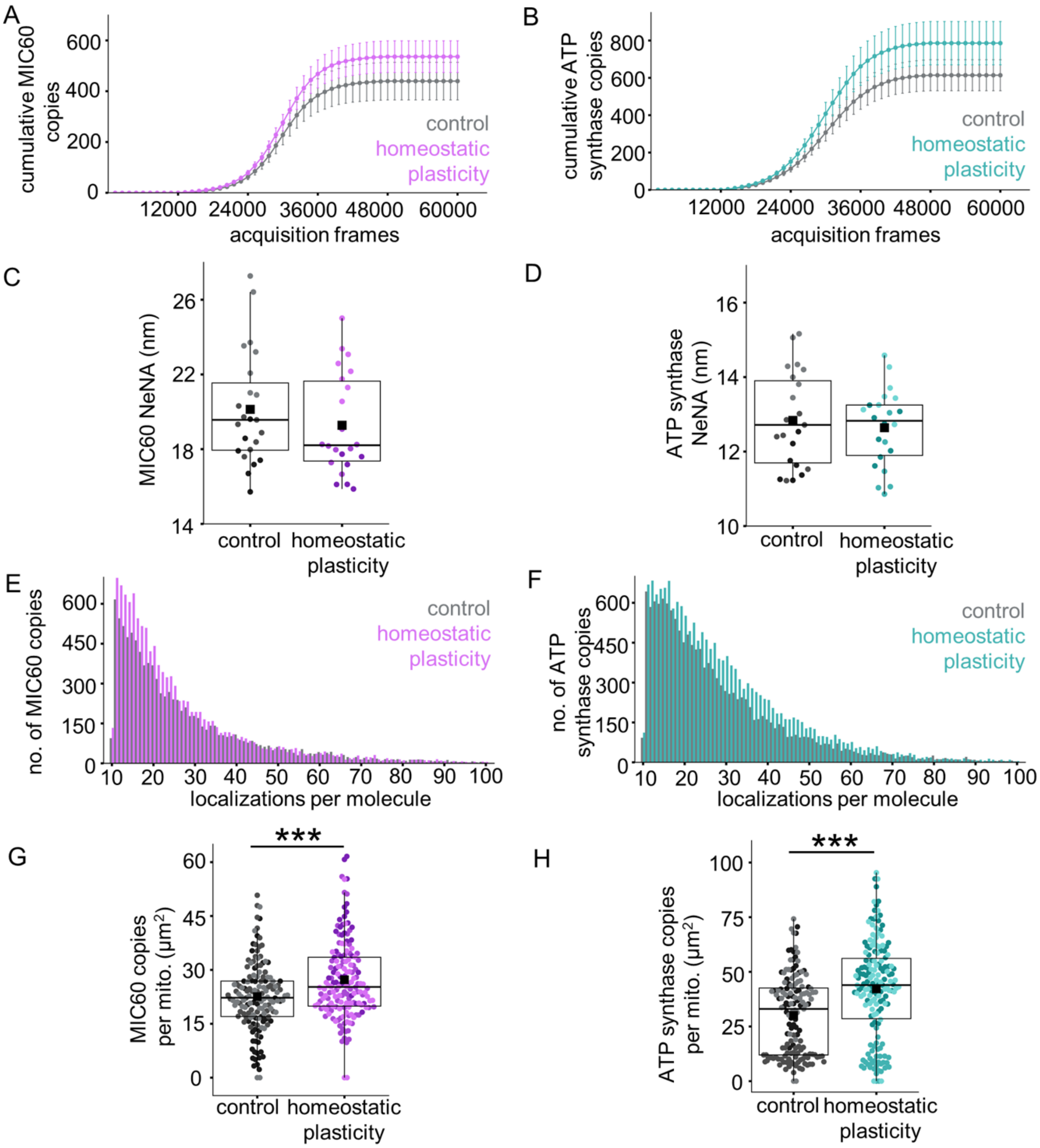

**Figure S9. DNA-PAINT validation and quantification of MIC60 and ATP synthase**

**A, B** Cumulative detected copies for MIC60 (**A**, purple) and ATP synthase (**B**, turquoise) reach saturation over the acquisition in homeostatic plasticity-induced neurons and in respective controls (gray), confirming complete detection of all labeled molecules. n in neurons, biol. replicates: 23, 3 (MIC60, control), 22, 3 (MIC60, homeostatic plasticity), 23, 3 (ATP synthase, control), 22, 3 (ATP synthase, homeostatic plasticity). **C**, **D** Localization precision, estimated by nearest-neighbor analysis (NeNA) value, for MIC60 (**C**, purple) and ATP synthase (**D**, turquoise) in homeostatic plasticity-induced neurons and respective controls (gray). Different shades within each condition denote independent biological replicates. n in neurons, biol. replicates: 23, 3 (MIC60, control), 22, 3 (MIC60, homeostatic plasticity), 23, 3 (ATP synthase, control), 22, 3 (ATP synthase, homeostatic plasticity). **E**, **F** Distribution of localizations per copy assigned by the clustering algorithm for MIC60 (**E**, purple) and ATP synthase (**F**, turquoise) in homeostatic plasticity-induced neurons and respective controls (gray). n in copies, biol. replicates: 10114, 3 (MIC60, control), 11795, 3 (MIC60, homeostatic plasticity), 14121, 3 (ATP synthase, control), 17290, 3 (ATP synthase, homeostatic plasticity). **G**, **H** Average number of MIC60 (**G**, purple) and ATP synthase (**H**, turquoise) copies per mitochondrion increases following homeostatic plasticity relative to respective controls (gray). n in mitochondria, biol. replicates: 149, 3 (MIC60, control), 165, 3 (MIC60, homeostatic plasticity), 149, 3 (ATP synthase, control), 165, 3 (ATP synthase, homeostatic plasticity). Two-way ANOVA (condition*biol. replicates), Tukey test, p-values: $<0.0001$ (MIC60), $<0.0001$ (ATP synthase).

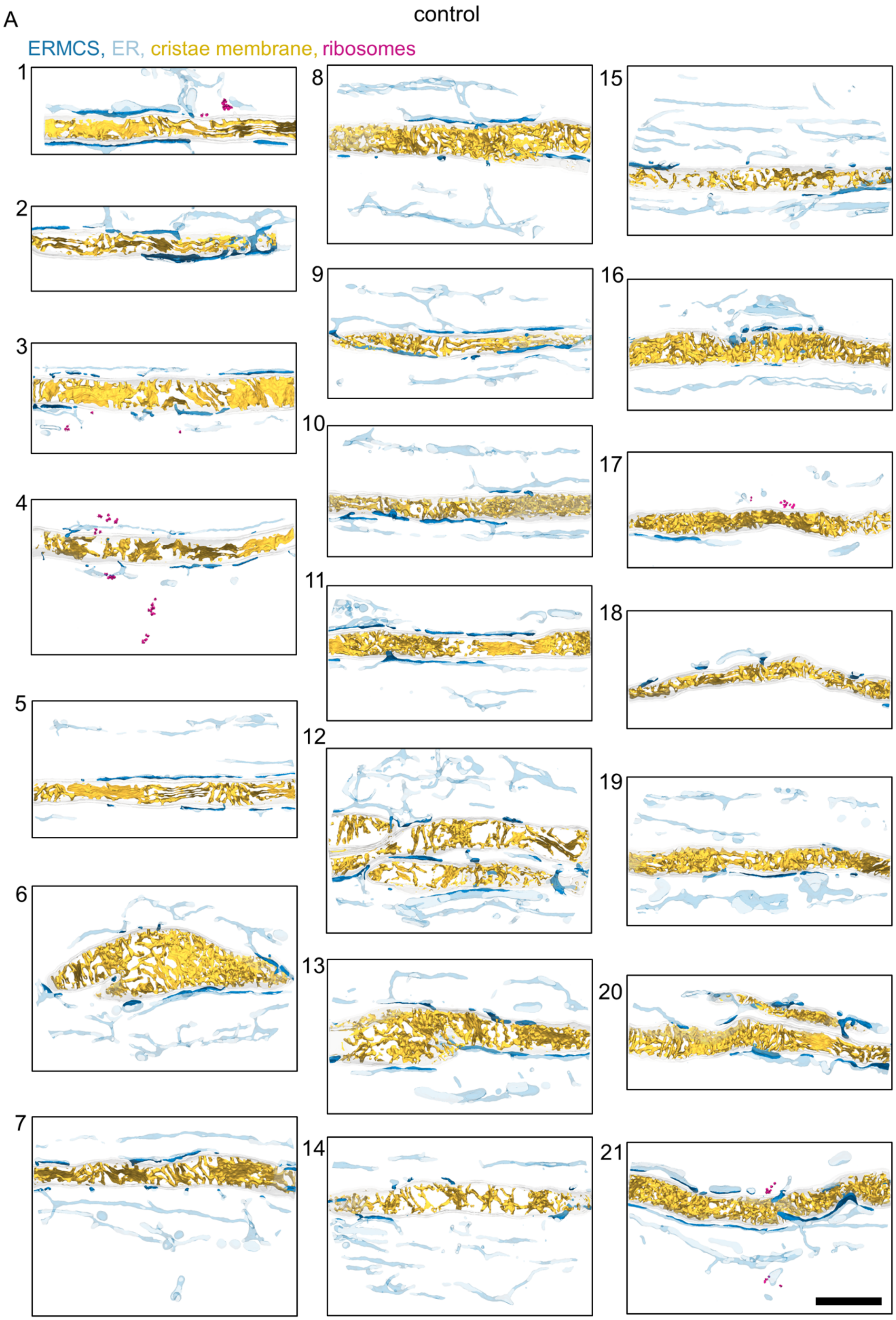
A
control
ERMCS, ER, cristae membrane, ribosomes

162

**Figure S10. Reconstructed electron tomograms showing MERCS and ribosomes in control.**

Reconstructed electron tomograms of juxtaspinal mitochondrial regions in control neurons (related to **Fig. 5B-E**), showing MERCS (dark blue), total ER (light blue), cristae membrane (yellow), and ribosomes (pink). n in reconstructions, biol. replicates: 21, 3. Mitochondrion 4 is the same as in **Fig. 5B**. Scale bar: 0.5 µm.

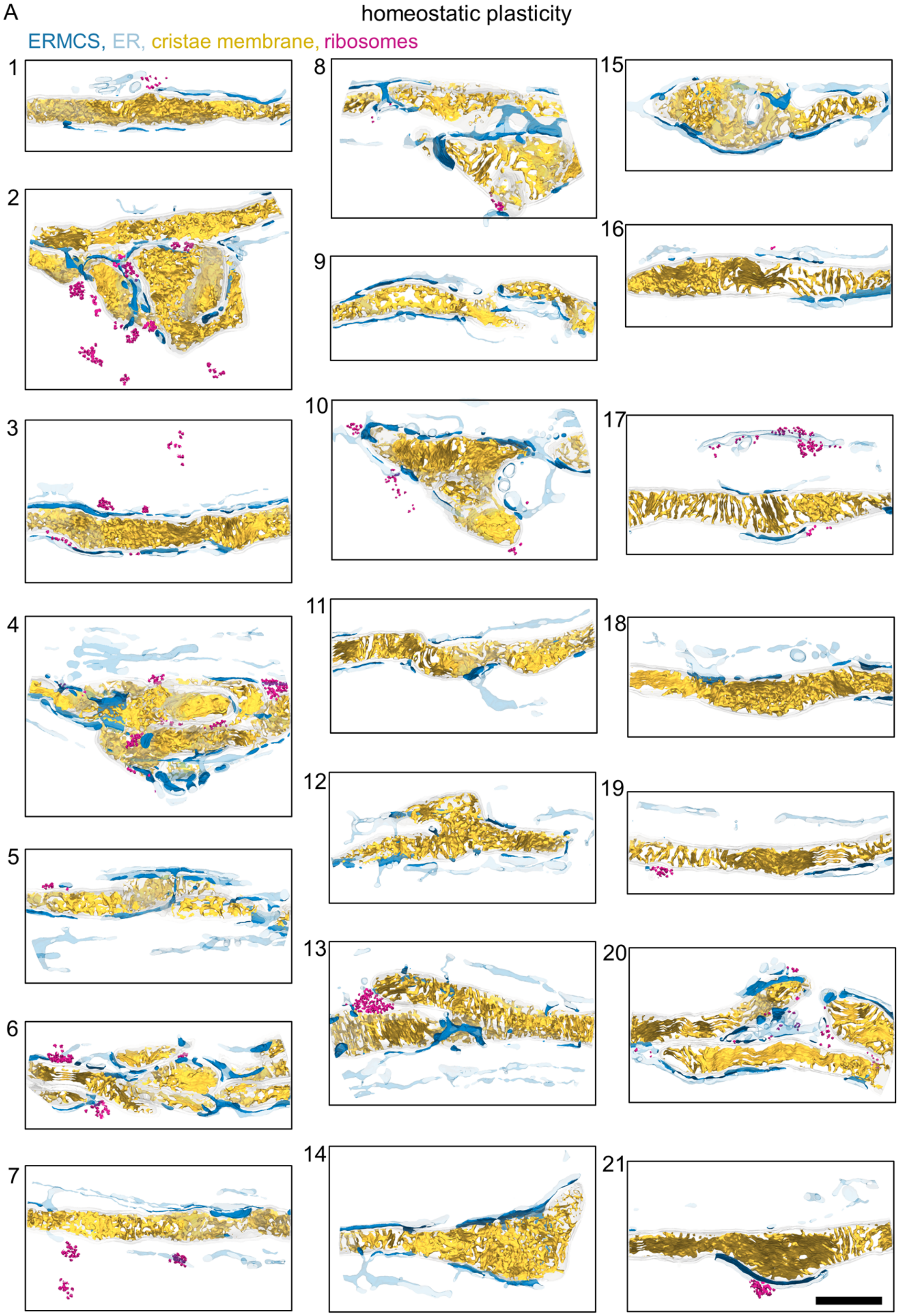
A
homeostatic plasticity
ERMCS, ER, cristae membrane, ribosomes

169

**Figure S11. Reconstructed electron tomograms showing increased MERCS and mitochondria-proximal ribosomes following synaptic plasticity.**

Reconstructed electron tomograms of juxtaspinal mitochondrial regions following homeostatic plasticity (related to **Fig. 5B-E**), showing MERCS (dark blue), total ER (light blue), cristae membrane (yellow), and ribosomes (pink). n in reconstructions, biol. replicates: 21, 3. Mitochondrion 2 is the same as in **Fig. 5B**. Scale bar: 0.5 µm.

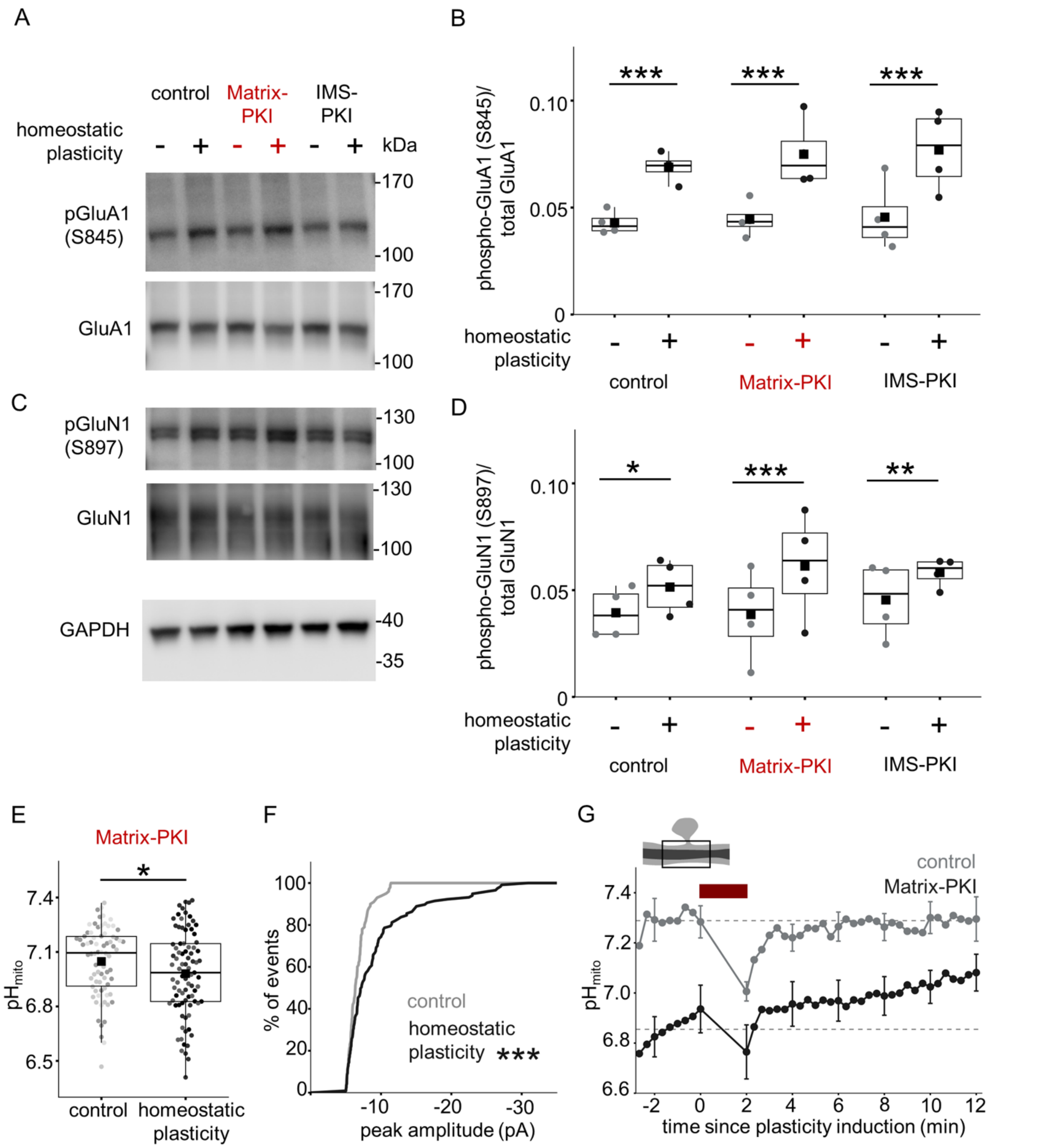

**Figure S12. Control experiments related to compartment-specific PKA inhibition**

**A, C** Representative Western blots measuring GluA1 (**A**) and GluN1 (**C**) phosphorylation following homeostatic plasticity induction in control (black), Matrix-PKI (red), and IMS-PKI (black) expressing neurons. GAPDH served as the loading control. **B, D** Average ratio of phosphorylated to total protein signal, showing increased GluA1 (**B**) and GluN1 (**D**) phosphorylation following homeostatic plasticity in control, Matrix-PKI-, and IMS-PKI-expressing neurons. n in blots, biol. replicates: 4, 4; 4, 4 (GluA1; GluN1: control, +/-homeostatic plasticity), 4, 4; 4, 4 (GluA1; GluN1: Matrix-PKI, +/-homeostatic plasticity), 4, 4; 4, 4 (GluA1; GluN1: IMS-PKI, +/-homeostatic plasticity). Two-way (condition*treatment) matched ANOVA, Tukey test, p-values: 0.0009, 0.0133 (control: GluA1, GluN1), 0.0003, 0.0005 (Matrix-PKI: GluA1, GluN1), 0.0002, 0.0099 (IMS-PKI: GluA1, GluN1). **E** In Matrix-PKI-expressing neurons, average $pH_{mito}$ shows a small but significant decrease following homeostatic plasticity (dark gray) relative to control (light gray). Different shades within each condition denote independent biological replicates. n in mitochondria, biol. replicates: 74, 3 (control), 85, 3 (homeostatic plasticity). Two-way ANOVA (condition*biol. replicates), p-value: 0.0478. **F** In Matrix-PKI-expressing neurons, the cumulative distribution of mEPSC peak amplitude shows a significant increase following homeostatic plasticity (black) relative to control (gray). n in neurons, biol. replicates: 9, 3 (control), 11, 3 (homeostatic plasticity). Kolmogorov–Smirnov test, p-value: 0.0001. **G** Average time course of $pH_{mito}$ in juxtaspinal mitochondrial regions in control (gray) and Matrix-PKI-expressing (black) neurons upon single-spine plasticity induction (red bar, 0.5 Hz, 120s) used to pH-correct $ATP_{mito}$ in **Fig. 6G.** n in mitochondria, biol. replicates: 8, 3 (control), 7, 3 (Matrix-PKI).
